# Dynamic Construction of Reduced Representations in the Brain for Perceptual Decision Behavior

**DOI:** 10.1101/284158

**Authors:** Jiayu Zhan, Robin A. A. Ince, Nicola van Rijsbergen, Philippe G. Schyns

## Abstract

Current models propose that the brain uses a multi-layered architecture to reduce the high dimensional visual input to lower dimensional representations that support face, object and scene categorizations. However, understanding the brain mechanisms that support such information reduction for behavior remains challenging. We addressed the challenge using a novel information theoretic framework that quantifies the relationships between three key variables: single-trial information randomly sampled from an ambiguous scene, source-space MEG responses and perceptual decision behaviors. In each observer, behavioral analysis revealed the scene features that subtend their decisions. Independent source space analyses revealed the flow of these and other features in cortical activity. We show where (at the junction between occipital cortex and ventral regions), when (up until 170 ms post stimulus) and how (by separating task-relevant and irrelevant features) brain regions reduce the high-dimensional scene to construct task-relevant feature representations in the right fusiform gyrus that support decisions. Our results inform the occipito-temporal pathway mechanisms that reduce and select information to produce behavior.

---

Over the past decade, there has been extensive studies of the regions of the brain that support face, object and scene recognition using different methodologies and modalities of brain measurements (e.g. electrophysiology, E/MEG and fMRI), including across species. A converging set of results now suggests a hierarchically organized architecture of brain regions, spanning the occipital and temporal lobes (1–17), where categorizations unfold over the first few hundred milliseconds of post stimulus processing (18–22). This same architecture is flexibly involved in multiple categorization tasks that require multiple representational bases. With extensive knowledge of the *where* and *when*, the next challenge is to unravel the thorny *how*. That is, how detailed information processing mechanisms in the occipitoventral pathway dynamically implement flexible visual categorization by selecting, from the high-dimensional input, the low-dimensional information basis required for behavior? In other words, how does our brain extract, from the features of the visual scene that it initially represents, those that are actually useful for the categorization task?

To start addressing this considerable challenge, we used Dali’s ambiguous painting *Slave Market with Disappearing Bust of Voltaire* (see Figure 1, Stimulus) as a case-study stimulus because it is a complex visual scene that affords two distinct perceptions. We applied the Bubbles technique (Gosselin & Schyns, 20001) to characterize the specific features of the visual scene associated with each perceptual decision. Specifically, on each experimental trial, we randomly sampled information from the scene (see Figure 1, Stimulus Sampling, for an illustration) while simultaneously measuring perceptual decisions (perceiving “the nuns,” “Voltaire” or “don’t know”) and the dynamic brain activity on 12,773 MEG voxels between 0 and 400 ms post-stimulus (see *Methods, Observers, Stimuli, Procedure, MEG Data Acquisition*).

**Figure 1.**
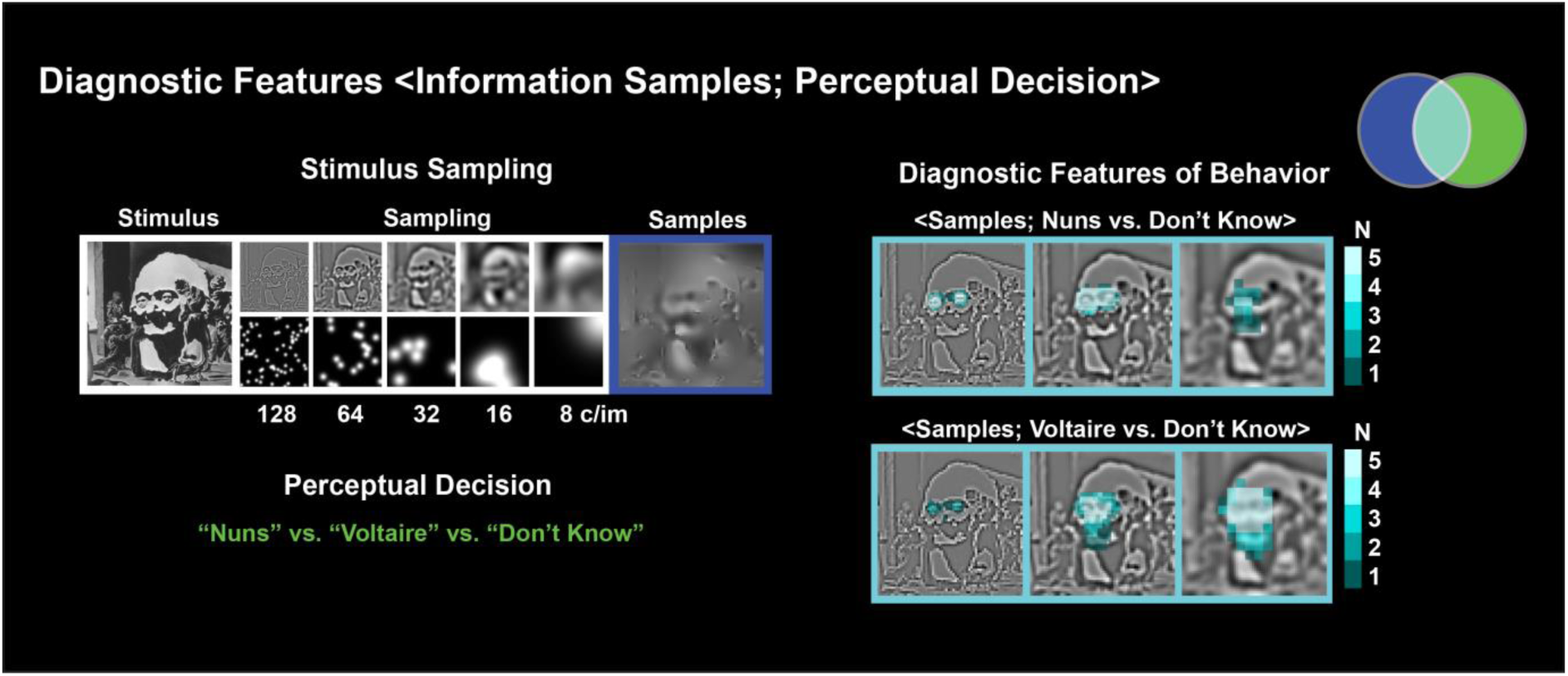
Experimental Design and Diagnostic Features. On each experimental trial, we decomposed the original stimulus into 6 spatial frequency (SF) bands (band 6 not shown) of one non-overlapping octave each, starting at 128 cycles per image (c/im). Using the Bubbles procedure, we sampled visual information independently in each SF band with randomly positioned Gaussian apertures. We added these samples across SF bands to the constant 6^th^ SF band to generate one experimental stimulus (highlighted with a dark blue frame), from which we recorded the observer’s perceptual decision behavior (i.e. perceiving “the nuns,” “Voltaire,” “don’t know”—the samples shown typically elicited “Voltaire”). With Mutual Information, in each observer we quantified the single-trial relationship, for each sampled pixel in the 5 SF bands and separately for trials with responses <Information Samples; “nuns” vs. “don’t know”> and <Information Samples; “Voltaire” vs. “don’t know”>. The cyan framed images report the number of observers with significant pixels (FWER *p* < 0.01, FWER corrected) in the first three SF bands, revealing the features most diagnostic for responding “the nuns” (the two small faces in SF band 1) and “Voltaire” (the broad face in SF band 3).

Across trials, we performed a multi-level analysis of several relationships of the single-trial data triple <Information Samples; MEG Voxel Activity; Perceptual Decision> within a novel information theoretic framework. To preview, we first coupled sampled stimulus information with behavior to reveal the stimulus features that subtend perceptual decision. We then independently coupled sampled information with MEG voxel amplitudes to characterize the stimulus features the brain dynamically represents. In our figures, we schematize the relationships of the data triple with Venn set diagrams to progressively introduce the different levels of our analysis framework and to present its results. The outcome is a novel interpretation of occipito-ventral stream activity in terms of the dynamic representation of the high-dimensional input, including the reduction of behaviorally irrelevant information and the selection of the lower dimensional basis that supports behavior.

## RESULTS

### Diagnostic features of behavior: <Information Samples; Perceptual Decision>

First, we addressed the behavioral level and identified the low-dimensional information basis of perceptual decision. For each individual observer, we evaluated the relationship between information randomly sampled from the painting on each trial (the variable represented with a blue set in Figure 1) and the corresponding perceptual decisions (the variable represented with a green set in Figure 1). We represent the relationship between these two variables with the cyan Venn diagram intersection and call its contents the diagnostic features of behavior—i.e. the stimulus features that support each observer’s perceptual decisions (23–26). To extract diagnostic features, we used Mutual Information (MI, (27)), to quantify the relationship (Family-wise error rate (FWER) *p* < 0.01, one-tailed) between the presence of each image pixel determined by random bubble samples on each trial and the observer’s varying perceptual decision. We computed this relationship separately for the behavioral contrasts <Information Samples; “the nuns,” vs. “don’t know”>, excluding “Voltaire” trials and <Information Samples; “Voltaire”, vs. “don’t know”>, excluding “nuns” trials (see *Methods, Diagnostic Features of Behavior*).

Figure 1 presents the diagnostic features of behavior (see Supplementary Figure 1 for those of each individual observer). As shown with increased brightness, all observers (N = 5) used the local left and right nun’s faces at higher spatial frequencies (HSF) to respond “the nuns,” whereas all observers used the global face of Voltaire at lower spatial frequencies (HSF) to respond “Voltaire” (see the cyan framed images in Figure 1). Since diagnostic features influence behavior, the output of the brain, we know that the observer’s brain must represent at least these features. That is, when we observe differential neural activity in the occipitoventral pathway in response to different visual categories (19, 28–30), diagnostic features likely influence this activity between stimulus onset and observer decision. Next, we show that the MEG signals do indeed represent all diagnostic features over time.

### Representation of Features in the Brain: <Information Samples; MEG Voxel Activity>

We switched to the brain to identify the features (diagnostic or not) that MEG activity represents. With a data-driven analysis, for each observer we quantified with MI (27) the single trial relationship <Information Samples; MEG Voxel Activity> to first extract and then to precisely quantify the features that their brain represents (see *Methods, Representation of Features in the Brain*). These analyses produced a 3D matrix in which, for each feature represented in the observer’s brain (1^st^ dimension), MI values indicate the effect sizes of the feature representation (FWER *p* < 0.05, one-tailed) over 12,773 MEG voxels (2^nd^ dimension), every 2 ms between 0 and 400 ms post-stimulus (3^rd^ dimension)—i.e. a brain feature-by-voxel-by-time 3D matrix per observer. These per-observer representation matrices form the bases of all remaining analyses. They are unique to our approach because, as we will see, they enable more direct information processing interpretations of brain activity at a fine granularity of task features.

Figure 2a presents the resulting color-coded brain features consistently represented across observers and projected onto a common basis for display purposes (see *Methods, K-means of Brain Features* and Supplementary Figure 2 for the brain features of each individual observer). Even casual inspection of Figure 2a reveals that some brain features display the same visual information as the diagnostic features of behavior (i.e. the local nun’s faces at higher spatial frequencies and the global face of Voltaire at lower spatial frequencies) whereas others do not (e.g. the dark brown feature).

**Figure 2.**
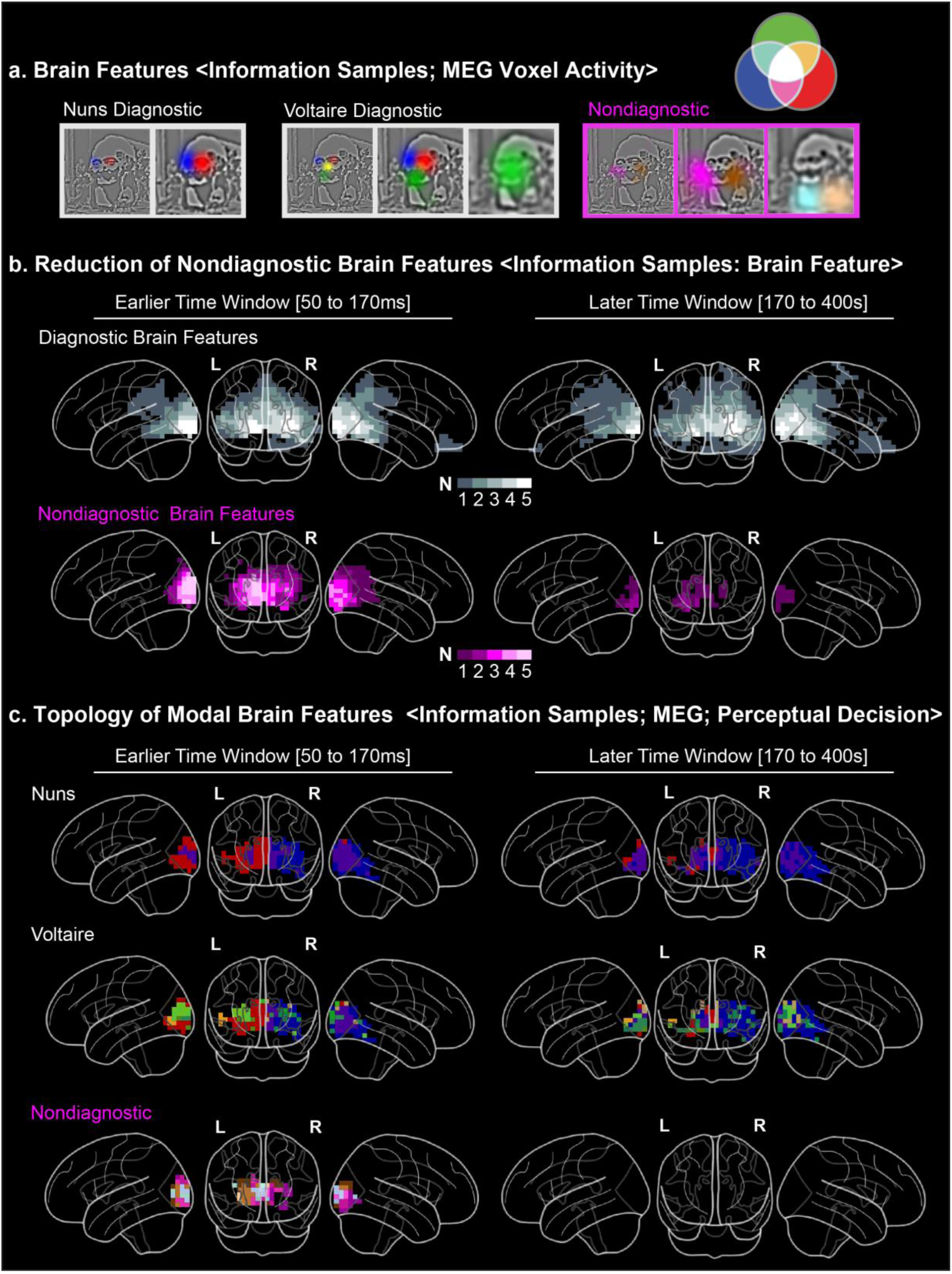
Dynamic Representation of Diagnostic and Nondiagnostic Brain Features. *a. Brain Features.* White frames highlight the diagnostic features that MEG voxels represent, separately presented for perceptual decisions of “the nuns” and “Voltaire.” Magenta frames highlight the features that MEG voxels also represent though they are not diagnostic of the perceptual decisions. The brain features are represented in a common feature basis resulting from a k-means analysis performed across the brain features of individual observers for display purposes. *b. Reduction of Nondiagnostic Brain Features.* The white (vs. magenta) glass brains report the voxels that represent at least one significant (FWER *p* < 0.05, one-tailed) diagnostic (vs. nondiagnostic) brain feature in an earlier (vs. later) time window post stimulus, where shading denotes the number of observers for whom these criteria hold true. Comparison of the earlier and later time windows of representation of diagnostic and nondiagnostic brain features indicates, for all observers, a consistent reduction of the latter, post 170 ms, at the juncture between the occipital and ventral regions. *b. Topology of Modal Brain Features.* In the earlier and later time windows, glass brains present the color-coded diagnostic brain features represented across a majority of observers for each voxel. Note the expected topological representation in occipital cortex and also that only diagnostic features are represented deep in ventral cortex post 170 ms.

In the Venn diagram (see Figure 2a), the white triple set intersection denotes visual scene information that influences both behavioral and brain measures (i.e. diagnostic brain features), whereas the magenta intersection denotes scene information that influences brain measures but not behavior (i.e. nondiagnostic brain features). We divided the brain features accordingly, into diagnostic and nondiagnostic for the decision task. Then, for each diagnostic feature, we established the specific decision it was primarily associated with (i.e. perceiving “the nuns” vs. “Voltaire”, see *Methods, Diagnostic and Nondiagnostic Brain Features*).

### Stimulus Representation Reduction between the Occipital and Ventral Cortex around 170 ms post stimulus

The glass brains in Figure 2b summarize the representation of diagnostic and nondiagnostic brain features over two time windows—i.e. [50-170 ms] and [170-400 ms] post stimulus flanking the important N/M170 event associated with visual categorizations (21, 31). Voxel brightness indicates the number of observers with statistically significant (FWER *p* < 0.05, one-tailed) representation of at least one diagnostic brain feature (white glass brains) or one nondiagnostic brain feature (magenta glass brains) in each time window, see *Methods, Flow of Brain Features*.

The glass brains reveal a consistent pattern across observers that can be interpreted as a reduction of the representation of stimulus information in the brain. Whereas voxels in all occipito-ventral regions represent diagnostic brain features over the first 400 ms of processing, nondiagnostic brain features are primarily represented in occipital cortex, and only during the first 170 ms of processing. Following this, nondiagnostic features extinguish across the brain of each observer (compare the white and magenta glass brains across the Earlier and Later time windows in Figure 2b). Figure 2c complements these data with a color-coding that shows, voxel by voxel, the specific diagnostic and nondiagnostic brain features represented across a majority of observers in each time window (see Methods, *Topology of Brain Feature Representations*).

In the earlier time window, feature representation reflects the expected topological projection in early visual cortex. Within the resolution of MEG voxels, we can observe a contral-lateral representation of the diagnostic left HSF nun’s face (color-coded in blue) and the right nun’s face (color-coded in red) and an upside-down contra-lateral representation of the eyes of Voltaire (color-coded in blue and red) in relation to the more broadly and bilaterally represented LSF Voltaire face (color-coded in green). As expected, the brown and purple nondiagnostic features flanking the centre of the painting are also contra-laterally coded (see Nondiagnostic Features glass brains in Figure 2c). However, note that only diagnostic features are represented deep into the right fusiform gyrus in both time windows (see also Supplementary Figure 2 that develops these findings for the maximally represented feature in the 3D feature-by-voxel-by-time matrix of each observer).

In sum, representation of diagnostic and nondiagnostic brain features in each observer reveals a consistent pattern of data reduction at a spatio-temporal junction located between the occipital and ventral regions, 170 ms post stimulus. Following this junction, the occipito-ventral pathway only represents diagnostic brain features. We now detail the information processing that happens before and after the junction: First, the pre-170 ms dynamic reduction of non-diagnostic brain features in occipital cortex; then, a post-170 ms right fusiform gyrus mechanism that dynamically constructs distributed representations of diagnostic brain features for perceptual decisions.

### Dynamic Reduction of Nondiagnostic Brain Features in the Occipito-Ventral Pathway

In each observer, we now demonstrate that nondiagnostic and diagnostic feature representations travel across occipital cortex towards the ventro-dorsal junction (i.e. ITG, MTG, STG and FG) like two wavefronts that diverge around 170 ms post stimulus. To this end, we used each observer’s 3D feature-by-voxel-by-time representation matrix to compute a curve of maximum representation (i.e. MI value) of nondiagnostic and diagnostic features, separately (see Methods, *Diagnostic and Nondiagnostic Feature Representation in Occipital Cortex*). The resulting representation curves are plotted in Figure 3a and 3b for one typical observer. In each plot, we ranked and color-coded each curve by its onset time, where cyan indicates earlier onsets and yellow indicates later onsets. Brain scatters adjacent to the plots use the same color-code to locate the voxels associated with the curves.

**Figure 3.**
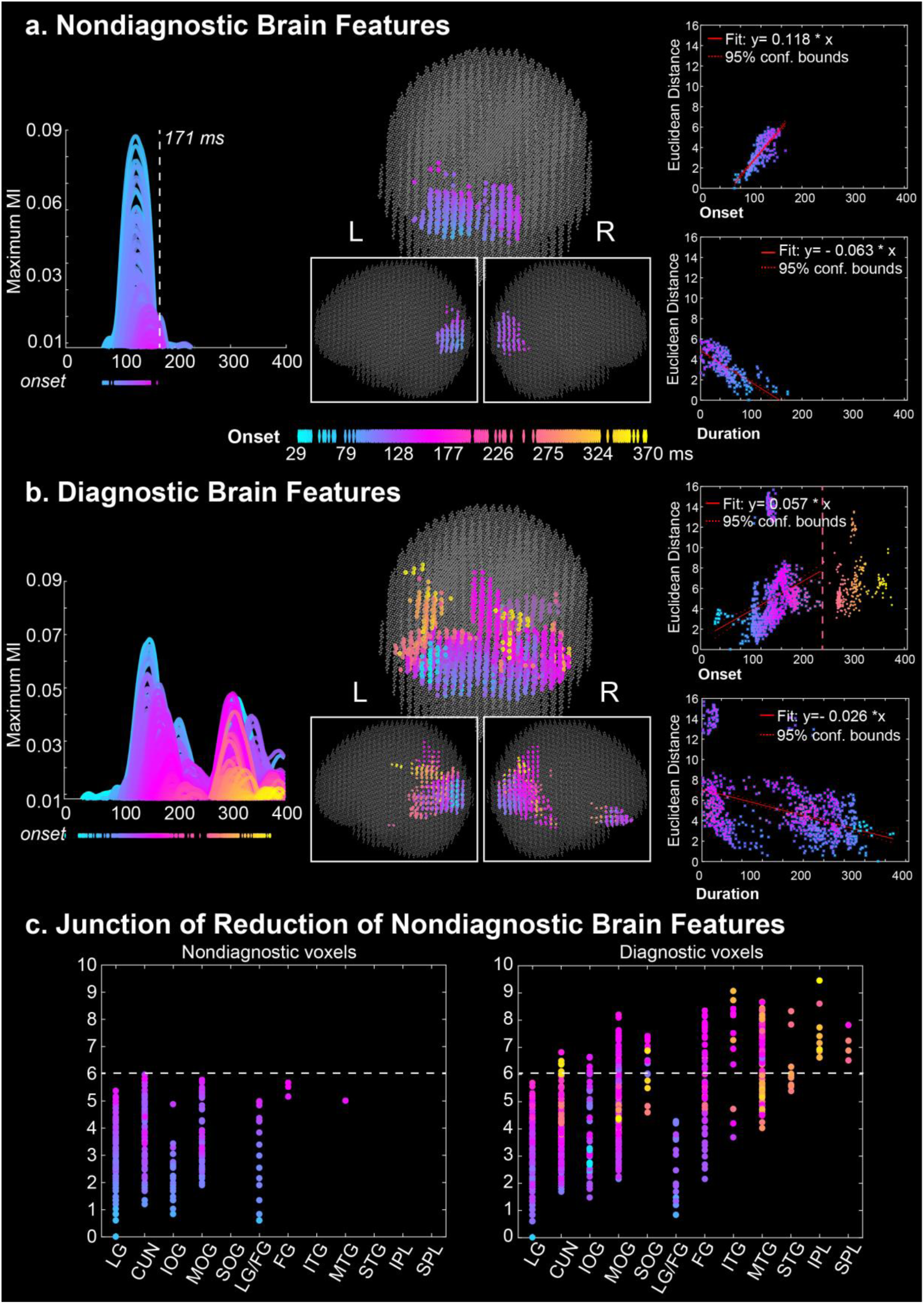
Reduction of Nondiagnostic Brain Feature Representations in Occipital Cortex in One Typical Observer. *a. Nondiagnostic Brain Feature.* <u>Left panel:</u> Maximum nondiagnostic brain feature representation curves for each voxel that represented at least one nondiagnostic brain feature, color-coded by ranked onset time (blue, early; yellow, late). Onset times are projected underneath the plot. The vertical dashed line represents the maximum offset time of the furthest voxels representing nondiagnostic features (i.e. highest Euclidean distance to the cyan voxel of earliest onset). <u>Middle panel:</u> To locate the voxels contributing curves in the left panel, the color code of onset time is used in 3 views of a 3D brain scatter. <u>Right panel</u>. The top scatter shows a linear relation between voxel onset times and their Eucidean distances to the initial onset voxel demonstrates. The bottom scatter shows a linear decrease of duration of nondiagnostic feature representation on voxels with their increasing distance from the initial onset voxel. *b. Diagnostic Brain Features.* Same caption as in panel a for diagnostic brain features, with later onsets (pink to yellow colors) in ventral and dorsal regions. *c. Junction of Reduction of Nondiagnostic Brain Features.* <u>Left panel</u>. Voxels color-coded by onset times are pooled by anatomical brain region (X axis) and scattered according to their Euclidean distance to the initial onset voxel of nondiagnostic representation on the Y axis. <u>Right panel</u>. Same caption for diagnostic voxels. Horizontal dash line indicates the brain regions of furthest representation of nondiagnostic features. LG/FG on X axis comprises voxels located near to Lingual gryus (see Supplementary Figure 7 for location).

Figure 3a illustrates that nondiagnostic feature representations travel as a wavefront that reduces in duration as it progresses through the occipital cortex. The rightmost top inset of Figure 3a demonstrates the wavefront property by showing that the onset times of the corresponding voxels (X axis, range [60-160 ms]) increase with their Euclidean distance from the initial onset voxel (Y axis, range [0 – 6], Y = 0.118 * X – 3.546, robust linear regression, *p* < 0.001). The rightmost bottom inset of Figure 3A demonstrates the reduction of duration by showing that representation duration (X axis) reduces with voxel Euclidean distance from the initial onset voxel (Y axis, Y = - 0.063 * X + 4.906, robust linear regression *p* < 0.001). Together, these data confirm that the wavefront of nondiagnostic feature representations collapses as it travels into the occipital cortex. In contrast, the same computations applied to diagnostic features in Figure 3B confirm that the diagnostic wavefront progresses past 170 ms, with later onsets deeper into ventral and dorsal regions.

Finally, the two panels of Figure 3c identify the anatomical brain regions (X axis) where the nondiagnostic and diagnostic wavefronts diverge. In each region, dots represent voxels color-coded by their onset time. The Y axis shows the Euclidean distance of each voxel in relation to the initial onset voxel, separately for nondiagnostic (left panel of Figure 3c) and diagnostic representations (right panel of Figure 3c). The dashed white horizontal line of collapse reveals that the nondiagnostic wavefront in the left panel extends ventrally from its initial onset in lingual gyrus up to the junction with the temporal gyrus and FG, and dorsally with inferior and superior parietal lobes. In contrast, the diagnostic wavefront in the right panel extends further into ventral and dorsal regions (see pink to yellow colors in Figure 3B). See *Methods, Diagnostic and Nondiagnostic Feature Representation and Occipital Cortex*. Supplementary Figure 3-1 to 3-4 show similarly divergent wavefronts in the occipito-ventral pathway of each remaining observer.

**Figure 4.**
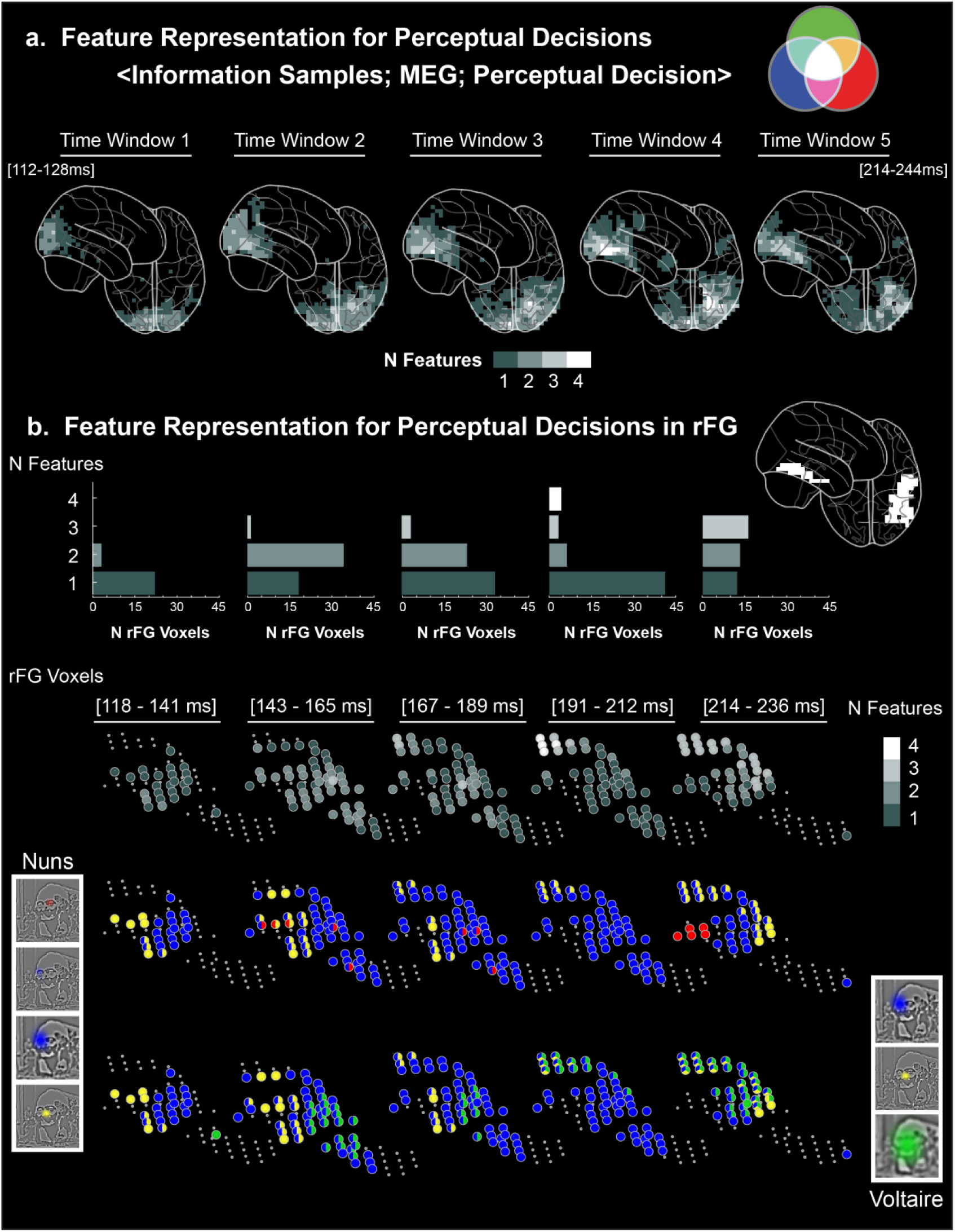
Dynamics of Perceptual Decision Representations in the rFG. *a. Feature Representation for Perceptual Decisions <Diagnostic Brain Features; MEG Voxel Activity; Perceptual Decision>.* Glass brains illustrate the evolution of the complexity of feature representations for behavior (i.e. redundant features) on brain voxels over 5 time windows. The grey level of a voxel corresponds to the median redundant feature complexity (i.e. the median number of different redundant features computed across 5 individual observers) *b. Feature Representations for Perceptual Decisions in rFG.* The top panel expands the evolution of redundant feature complexity for the rFG voxels of one typical observer. Grey levels indicate the number of different redundant features that a rFG voxel represents in each time window. Histograms underneath show the number of voxels that code 1 to 5 redundant features in each time window. In the bottom panels, color codes reveal the actual redundant features represented on each rFG voxel, for each perceptual decision, in each time window.

### Dynamic Construction of Representations for Behavior in the Right Fusiform Gyrus: <Information Samples; MEG Voxel Activity; Perceptual Decision>

We now turn to the fate of the diagnostic brain features whose representation progresses past the ventro-dorsal junction. A prevalent hypothesis is that visual information represented early and separately across the left and right occipital cortices (32) later converge into the right ventral pathway (33, 34), where lateralized representations become bilateral to support visual cognition such as perceptual decisions (35). However, conclusive testing of this hypothesis remains challenging for at least two reasons. First, ventral responses to perceptual categories are only understood when the determinants of the responses—i.e. the represented features driving ventral responses—are characterized. Second, even when features are characterized, it is necessary to demonstrate that they are represented specifically to support behavior.

To directly address these points and localize where, when and how the brain constructs representations for perceptual decisions, we directly quantified the triple white set intersection in Figure 4 with a novel information theoretic measure of redundancy—i.e. redundancy of <Brain Feature; MEG Voxel Activity; “the nuns,” “Voltaire,” “don’t know”>. Redundancy can precisely quantify representations of diagnostic features in the brain for behavior. We call these representations “redundant features.” In each observer, we computed the redundant features (FWER *p* < 0.05, one-tailed) for each brain voxel and time point between 112-244 ms post stimulus, the time period during which rFG represents the diagnostic features that pass the occipito-ventral junction--i.e. encompassing an extended N/M170 time course for visual feature integration (22, 36, 37), see *Methods, Feature Representation in the Brain for Perceptual Decision*. We split time into 5 windows and for each brain voxel estimated its representational complexity by summing the number of different redundant features that it represents within each time window. For example, a voxel representing Voltaire’s redundant green bust, left blue eye and yellow nose would have a complexity value of 3. Across observers, we then computed the median complexity of each voxel per time window and coded this number with a grey level in the glass brains of Figure 4a (where median complexity varied between 0 and 4) to reveal where and when the complexity of feature representations for perceptual decisions develops.

The glass brains show that representational complexity peaks across observers in voxels located at the top of the right fusiform gyrus (rFG, cf. 4^th^ time window, see Figure 4a). Specifically, representational complexity peaks in these voxels in the 4^th^ time window for 4/5 observers and in the 3^rd^ time window for the remaining observer—i.e. between 163ms and 218ms post stimulus. We now detail the evolution of complexity, in Figure 4b for one typical observer, and in Supplementary Figures 4-1 to 4-4 for each remaining observer. The histograms in Figure 4b indicate the number of rFG voxels (X axis) that represent 1 to 4 redundant features within each time window (Y axis) for this observer. The lower panels of Figure 4b characterize the dynamics of redundant features representations on rFG voxels to decide “the nuns” and “Voltaire,” leading to increasingly complex distributed representations in the top rFG voxels. For example, in the 4^th^ time window these voxels represent three redundant features to respond “Voltaire”—i.e. green bust, left blue eye, and yellow nose. Such dynamics of increasing representational complexity for perceptual decisions at the top of rFG was common to all observers (see Supplementary Figures 4-1 to 4-4).

In sum, we showed that the diagnostic features passing through the occipito-temporal junction into the rFG agglomerate over time at the top of the rFG over a few voxels where simultaneously represented redundant features form distributed representations of increasing complexity that support perceptual decisions.

## DISCUSSION

Here, we studied where, when, and how brain regions dynamically construct representations of a stimulus to deliver perceptual decision behaviors. To achieve this, we used a novel information theoretic framework that quantifies the intersections of single trial information sampling, MEG source signals, and perceptual decision. First, we coupled sampled stimulus information with perceptual decision to reveal the stimulus features that subtend behavior. We then independently coupled sampled information with MEG voxel amplitudes to characterize the stimulus features the brain dynamically represents. Using these data, we first showed that both diagnostic and nondiagnostic feature representations travel like two wavefronts in occipital cortex and which diverge around 170 ms post-stimulus. Specifically, the nondiagnostic wavefront collapses at the occipito-ventral junction around 170 ms, which thus reduces the stimulus information that was initially represented. However, the diagnostic wavefront progresses past this junction and into ventral and dorsal regions, which thus retains a reduced subset of the stimulus features. Second, using a new information theoretic measure that quantifies the white triple set intersection, we showed that the diagnostic features progressing past the occipito-ventral junction agglomerate over time on specific voxels at the top of the rFG. This agglomeration constructs the distributed feature representations of increasing feature complexity that subtend perceptual decision. We replicated the main results of (1) dynamic reduction of nondiagnostic features prior to 170 ms in occipital cortex and (2) construction of distributed, increasingly complex feature representations at top rFG voxels to support behavior in each observer independently.

### Data Reduction in the Occipito-Ventral Pathway

We have documented a data reduction in the brain that implies a process that evolves over time from a state of many dimensions of stimulus representation to a state of fewer dimensions of stimulus representation. However, this many-to-fewer state transition does not imply that the transition involves only feedforward processes. Instead, the different hierarchical layers of the occipito-ventral pathway likely communicate with each other using both feedforward and feedback signals to implement the data reduction over time. Interactive architectures of this type are similar to well-known network models that can resolve ambiguity between hierarchically organized representations (38, 39). We subscribe to such an interactive organization whereby the selection of diagnostic features from the visual stimulus probably involves predictions arising from memory, which propagate down the layers of the visual hierarchy and interact with the bottom-up flow of stimulus information (40–44). However, due to the arrow of time, although we can visualize the feedforward flow of stimulus representation by coupling information samples with subsequent brain responses we cannot yet visualize the representation of top-down predictions in the brain (though see (42) for visualizations of predictions from behavior). Nevertheless, as we will now discuss, this interactive architecture could be further documented by visualizing successive transformations of the stimulus representation over time.

Here, we trace the dynamic representation of, for example, the nun’s face (i.e. the high spatial frequency pixels representing this feature in the 2D image) from occipital cortex into the ventral pathway. However, it would be naïve to assume that the nun’s face is represented as such in any of these brain regions. Instead, Gabor patches, for example, would provide a better model of the stimulus feature representation in early visual cortex (45). Likewise, a model tolerant to changes in size, rotation, and illumination would better reflect established properties of higher-level representations in the ventral pathway. Therefore, to better understand the content of representations along the visual hierarchy, we would need to move from sampling pixels across SFs to sampling an explicit hierarchical generative model of visual information. At each level of the model, the information sampled would ideally mirror the information represented at each level of the visual hierarchy. At the top level of the generative hierarchy, models of 3D objects would comprise relevant object parts and their articulations, all of which would be tolerant to changes in size, rotation, and illumination. At the lower level of the generative hierarchy, the object’s parts would project onto Gabor filters at various resolutions to finally synthesize the 2D stimulus. In such a hierarchical generation, each level of the model would be an explicit hypothesis on the representation formats of each layer of the visual hierarchy in the brain. To enable a finer-grained testing and interpretation of explicit representations along the visual hierarchy as described here (i.e. including reduction and selection) could be achieved by filling the blue set of our framework with samples from such generative models. Designing these generative models of visual information remains, in our view, a necessary but considerable challenge to understanding complex visual representations (though see (46) for faces, (47) for simple objects and (48) for scenes).

### Time Course of Information Processing in the Occipito-Ventral Pathway

The reduction of nondiagnostic features and the progression of diagnostic features at the occipito-ventral junction around 170 ms strongly suggests coincidence with the N/M170 event related potentials. For example, N/M170 sources have been located around the STS, the fusiform gyrus, or both (49, 50), and intracranial data suggest the involvement of inferior occipital gyrus in the generation of the N/ M170. Therefore, these regions and the timing of the N/M170 (which peak around 170 ms post stimulus) are indeed similar to the spatio-temporal junction we have identified here. A further similarity is that the N/M170 also reflects a network that represents and transfers features across the hemispheres of the brain. For example, in a face detection task where the two eyes are diagnostic, we showed that the N/M170 first represents the contra-lateral eye followed by the ipsi-lateral eye, which has been transferred from the opposite hemisphere (18). This process occurs over a time window that flanks the N/M170 peak and which is analogous to the time course of the construction of complex feature representations for perceptual decisions shown here. Further studies should explicitly test at least the following three hypotheses. First, that the N170 reflects the moment when, as illustrated here, the wavefronts of nondiagnostic and diagnostic features diverge. This would be compatible with the recurrent finding that the largest N170 amplitudes are associated with face stimuli. Since faces are important sources of social information for humans, they could have intrinsic diagnosticity with their information systematically transferred into the rFG for further social signal processing. A second and related hypothesis is that, as shown here (see also (18)), the pre- and post-170 ms time course in the rFG could reflect two processing stages. Pre-170 ms, rFG could buffer visual information that arises first from the contra-lateral visual field, followed by ipsi-lateral visual field information that is transferred from the left occipital hemisphere; post-170 ms, rFG would integrate this buffered information across the two visual fields. This hypothesis could be tested using contrasting tasks where the integration of information in the rFG is strictly necessary for behavioral decisions (e.g. judging whether the left and right hemifields present the same or different information) vs. unnecessary (e.g. judging whether information appears in either hemifield). Finally, a third hypothesis should test whether the data reduction shown here reflects a specific process of figure ground separation in the image, or a more generic process of selection of feature with relative task-specific weights. This would entail using scenes with faces and objects and instruct observers to categorize either the faces or the objects and compute what information is reduced in each.

Our paper addresses an important question about the time course of the construction of feature representations for perceptual decisions. Our framework also offers a new approach to studying decision making by modulating the relevance (i.e. diagnosticity) of specific features for behavioral decision, and measuring how feature diagnosticity affects their accrual in the brain for perceptual decisions. We can also quantify the effect of each feature on perceptual decisions and measure how feature accrual induces the nonlinearity of decision making. Critically, we can explicitly model, rather than assume, the accrual process that is central to most models of decision making in humans. However, there remains an outstanding question as to whether perceptual decisions are primarily associated with the P300 (51), a positive event related potential arising in parietal regions (36), or whether decisions start earlier such as over the time course of the N170. Here, our quantification of the white triple set intersection with redundancy revealed the construction of decision-specific distributed representations with diagnostic features occurring in the top voxels of the rFG before 250 ms. While we can be sure that these rFG representations subtend perceptual decisions, it remains unclear whether actual decision making occurs there and then. Indeed, decision itself could be a process that is distributed across the construction of a diagnostic representation in the rFG and the elicitation of actions in a network spanning the fusiform gyri, parietal, and pre-motor cortices (52).

In sum, our information theoretic framework enables a more precise and principled study of information processing in the brain for behavior (53). In essence, this framework allows the characterization of diagnostic information, tracing the flow of its representation in brain activity between stimulus onset and perceptual decision, and contrasting it to the representation of nondiagnostic information. Here, we have demonstrated the strength and feasibility of this framework in understanding fundamental mechanisms of visual perception and categorization but it is also generalizable to other sensory modalities.

## METHOD

### Observers

Five observers with normal (or corrected to normal) vision participated in the experiment. We obtained informed consent from all observers and ethical approval from the University of Glasgow Faculty of Information and Mathematical Sciences Ethics Committee.

### Stimuli

We cropped the ambiguous portion of Dali’s *Slave Market with the Disappearing Bust of Voltaire* to retain the bust of Voltaire and the two nuns. Image size was 256 x 256 pixels and presented at 5.72° × 5.72° of visual angle on a projector screen. On each trial, we sampled information from the cropped image using bubble masks made of randomly placed Gaussian apertures to create a different sparse stimulus. We explain the sampling procedure below and refer the reader to Figure 1, Stimulus Sampling. We decomposed the Dali image into six independent Spatial Frequency (SF) bands of one octave each, with cut-offs at 128 (22.4), 64 (11.2), 32(5.6), 16 (2.8), 8 (1.4), 4 (0.7) cycles per image (cycles per degree of visual angle), respectively. For each of the first five SF bands, a bubble mask was generated from a number of randomly located Gaussian apertures (the bubbles), with a standard deviation of 0.13, 0.27, 0.54, 1.08, and 2.15 degrees of visual angle, respectively. We sampled the image content of each SF band by multiplying the bubble masks and underlying greyscale pixels at that SF band, summed the resulting pixel values across SF bands and added the constant 6^th^ SF band (see Figure 1, Stimulus Sampling, 6^th^ SF band not shown) to generate the actual stimulus image (See Figure 1, Samples). The total number of 60 Gaussian apertures on each trial remained constant throughout the task, ensuring that equivalent amounts of visual information was presented on each trial, at a level found previously to maintain “don’t know” responses at 25% of the total response number (54). Since the 6^th^ underlying SF image was constant across trials, we performed all analyses on the 5 bubble masks controlling visibility, but reported only the first three because they represented most of the information required for perceptual decisions. For analysis, we down-sampled (bilinear interpolation) the bubble masks to a resolution of 64 × 64 pixels to speed up computation.

### Procedure

Each trial started with a fixation cross displayed for 500 ms at the centre of the screen, immediately followed by a stimulus generated as explained above that remained until response. We instructed observers to maintain fixation during each trial, and to respond by pressing one of three keys ascribed to each response choice—i.e. “the nuns”, “Voltaire”, or “don’t know.” Stimuli were presented in runs of 150 trials, with randomized inter-trial intervals of 1.5–3.5s (mean 2s). Observers performed 4–5 runs in a single day session with short breaks between runs. Observers completed the experiment over 4–5 days.

### MEG Data Acquisition

We measured the observers’ MEG activity with a 248-magnetometer whole-head system (MAGNES 3600 WH, 4-D Neuroimaging) at a 508 Hz sampling rate. We performed analysis with the FieldTrip toolbox (55) and in-house MATLAB code, according to recommended guidelines (56). For each observer, we discarded runs based on outlying gradiometer positions in head-space coordinates. That is, we computed the Mahalanobis distance of each sensor position on each run from the distribution of positions of that sensor across all other runs. Runs with high average Malahanobis distance were considered outliers and removed. The distances were then computed again and the selection procedure was repeated until there were no outlier runs (Mahalonobis distances > 20). We high-passed filtered data at 1 Hz (4^th^ order two-pass Butterworth IIR filter), filtered for line noise (notch filter in frequency space) and de-noised via a PCA projection of the reference channels. We identified noisy channels, jumps and other signal artefacts using a combination of automated techniques and visual inspection. We then epoched the resulting data set (mean trials per observer 3396, range 2885–4154, see Supplementary Table 1) into trial windows (− 0.8s to 0.8s around stimulus onset) and decomposed using ICA, separately for each observer. We identified and projected out of the data the ICA sources corresponding to artefacts (eye movements, heartbeat; 3 to 4 components per observer).

We then low-pass filtered the data to 40Hz (3^rd^ order Butterworth IIR filter), specified our interest time period 0-400ms post stimulus, and performed the Linearly Constrained Minimum Variance Beamforming analysis (57) to obtain the source representation of the MEG data on a 6mm uniform grid. We low-pass filtered the resulting single trial voxel time courses with a cut-off of 25Hz (3^rd^ order Butterworth IIR filter, two-pass). In the following analysis, based on the obtained single-trial voxel activity time courses (12,773 MEG voxels, every 2ms between 0 - 400ms post stimulus), we analyzed the dynamic representation of features in the brain for perceptual decisions.

The following sections detail each step of the information processing pipeline. We refer the reader to Supplementary Figure 5 for a schematic graphic overview.

### Diagnostic Features of Behavior: <Information Samples; Perceptual Decision>

To compute the diagnostic features of perceptual decisions, we quantified the statistical dependence between the pair <Information Samples; Perceptual Decision> using Mutual Information (MI,(27)). On each trial, 5 real-valued Gaussian bubble masks multiply the visual information represented in 5 SF bands (see Figure 1, Stimulus Sampling, for an illustration). Thus, on a given trial, a real value represents the visibility of that pixel under a Gaussian bubble, with 1 indicating full visibility and 0 indicating no visibility. For each pixel of the bubble mask, we converted its random distribution of real values across trials into 2 bins We thresholded the bubble masks with a value of 0.2, with values below threshold representing “no to low visibility” and values above representing “low to full visibility”. We then used MI to quantify the statistical dependence between the binarized pixel visibility values and the corresponding observer responses, grouping “the nuns” vs. “don’t know” responses together in one computation (i.e. <Information Samples; “the nuns,” “don’t know”>) and the “Voltaire” vs. “don’t know” responses in the other (i.e. <Information Samples; “Voltaire,” “don’t know”>). These computations resulted in two MI perceptual decision pixel images per observer (see Supplementary Figure 1). To determine the statistical significance of MI pixels and address the problem of multiple comparisons, we used the method of maximum statistics (58). Specifically, for each of 10,000 permutations, we randomly shuffled the observer’s choice responses across trials, repeated the computation of MI for each pixel as explained and extracted the maximum MI across all pixels over the 5 SF bands. We used the 99^th^ percentile of the distribution of maxes across 10,000 permutations to determine the above chance significance of each MI pixel (FWER *p* < 0.01, one-tailed). Across observers, we reported the diagnostic pixels with significant MI in the first 3 SF bands that illustrate the consistency of the main diagnostic features underlying perceptual decision behaviors (see Figure 1, Diagnostic Features).

### Representation of Features in the Brain: <Information Samples; MEG Voxel Activity>

We measured single trial MEG activity (i.e. the bivariate amplitude and instantaneous MEG gradiant (35) on 12,773 sources, every 2 ms between 0 and 400 ms post stimulus. A high-dimensional 12,773 x 200 voxel-by-time matrix therefore structures the MEG data. We aimed to quantify, for each observer, the features of the stimulus that each cell of this matrix represents, if any. For each individual observer, we proceeded in three steps. First, we used MI to quantify in a reduced space of MEG activity (i.e. 60 ICA MEG sources x 75 8 ms time points) the relationship <Information Samples; MEG Activity> to derive with MI pixel images the features of the stimulus that the MEG activity represents over time. Second, with Non-negative Matrix Factorization (NMF) we reduced the MI pixel images across 60 MEG sources and 75 time points to reveal the main features that the brain represents. Finally, we quantified with MI the representation of each one of these features in the voxel-by-time MEG activity matrix (12,773 voxels x 200 time points), to derive a 3D feature-by-voxel-by-time matrix of MI values. We now detail the computations involved in each step. Throughout, we performed information calculations on continuous variables using the Gausian-Copula Mutual Information (GCMI) estimator (27). This estimator is a rank-based approach that gives a lower bound estimate of the true mutual information by quantifying Gaussian copula dependence between pairs of variables, but without any assumption on the marginal distributions of each variable.

#### Step 1. Computation of the Relationship <Information Samples; MEG Activity>

Though we aim to identify the features represented in each cell of the full voxel-by-time matrix of MEG activity, it is computationally impractical to directly compute the features from the single trial relationship <Information Samples; MEG Voxel Activity> due to the enormous dimensionality of the space—64 x 64 x 5 SF bands pixels x 12,773 voxels x 200 time points. Instead, we used the method of (35) which computes the relationship over the more computationally tractable matrix of 60 Independent Component Analysis (ICA) sources representing MEG activity over 75 time points that span 0 to 600 ms post stimulus every 8 ms. It also down-samples the stimulus sampling space to 64 pixels x 64 pixels x 5 SF bands, to evaluate fewer individual pixels in total.

#### Step 2: Computation of Brain Features

For each observer, the reduced matrix computed above comprised MI images in each cell, for a total of 4,500 MEG-pixel information images across 5 SF bands (60 ICA sources x 75 time points). We vectorized each (64 x 64 x 5 = 200,480) MEG MI image as a 20,480-dimensional vector. We applied Non-negative Matrix Factorization (NMF, (59)) to the set of 4,500 vectorized MEG MI images to characterize the main NMF features of the stimulus that modulate MEG source activity, resulting in 21–25 components per observer. We thresholded these NMF features by setting to zero the pixels with low MI values (< 15% of the maximum pixel value across SFs). We then normalized the NMF features (L2-norm). Henceforth, we call “brain features” the normalized NMF features of each observer that modulate the MEG activity of their brain.

#### Step 3: Computation of the Relationship < Brain Feature; MEG Voxel Activity> in the Full Voxel-by-Time MEG Activity Matrix

Here, we used the brain features computed above from the reduced matrix of ICA MEG sources to quantify their representation into each cell of the full voxel-by-time matrix. To this aim, first we computed the visibility of each brain feature into the information samples (i.e. bubble mask) presented as stimulus on each trial. That is, we spatially filtered (i.e. dot product) the bubble mask for that trial with the brain feature computed above, thereby producing a scalar value indicating the visibility of this feature on this trial. We call these real values “brain feature coefficients.” Next, for each cell of the full voxel-by-time MEG activity matrix, for each brain feature independently, we quantified the relationship <Brain Feature Coefficients; MEG Voxel Activity>, using GCMI.

For each brain feature, the above computations produced a voxel-by-time MI matrix that quantifies with effect sizes the relationship between each feature and MEG activity, at each voxel and time point—i.e. a time series that quantifies the representation of the feature. For each feature, we determined the statistical significance of each cell in its voxel-by-time matrix using a permutation approach and the method of maximum statistic to address multiple comparisons. Specifically, for each of 200 permutations, we randomly shuffled the brain feature coefficients values across trials and recalculated the MI of the single trial relationship <Randomized Brain Feature Coefficients; MEG Voxel Activity>, for each cell of the voxel-by-time matrix. In each cell, we computed the maximum of the resulting MI matrix for each permutation and used the 95^th^ percentile of this value across permutations as statistical threshold (i.e. FWER *p* < 0.05, one-tailed). This produced, for each observer, a 3D feature-by-voxel-by-time MI matrix used in the remaining analyses.

### K-means of Brain Features

As shown in Supplementary Figure 2, the different observers represented similar brain features in the task. This warranted the projection of the individual observer’s brain features onto a common basis to compare their dynamic representation in the brain. To this aim, we applied k-means clustering analysis by setting *k*, the number of clusters, to 25, to align the number of means to the maximum number of NMF brain features computed in any observer. We pooled the normalized NMF brain features of all observers, resulting in a 115 x 20480 matrix (115 NMF components in total for 5 observers, 64 pixels * 64 pixels * 5 SFs weights). We applied k-means (cosine similarity, 1000 repetitions) to this matrix (see Supplementary Figure 6 for the resulting k-means clusters). It is important to note that we performed all analyses on the specific brain features of each observer. We only indexed these individual features onto the common k-mean feature basis to report group results (e.g. in Figures 2 and 4).

### Diagnostic and Nondiagnostic Brain Features

For each observer, we formalized their diagnostic vs. nondiagnostic status of brain features by quantifying the single-trial MI relationship <Brain Feature Coefficient; “the nuns,” “Voltaire,” “don’t know”>. A strong relationship (i.e. MI above 75^th^ percentile of the distribution of MI across all brain features) would classify this brain feature as also diagnostic of perceptual decisions; a weak relationship (MI below 25^th^ percentile of the MI distribution) would classify this brain feature as nondiagnostic of perceptual decisions. Furthermore, we computed the importance of each diagnostic brain feature for perceptual decisions of “the nuns” (or “Voltaire”) by recomputing MI as just described, but using only the trials associated with responding “the nuns” vs. “don’t know” (or “Voltaire” vs. “don’t know”)—i.e. by recomputing <Brain Feature Coefficient; “the nuns,” “don’t know”> and <Brain Feature Coefficient; “Voltaire,” “don’t know”> independently. For each diagnostic brain feature, we determined it was diagnostic to perceiving “the nuns” (or “Voltaire”) when the perception-specific MI <Brain Feature Coefficient; “the nuns,” “don’t know”> (or <Brain Feature Coefficient; “Voltaire,” “don’t know”>) of the diagnostic brain feature was above the 75^th^ percentile of the distribution of the perception-specific MIs across all brain features (see Figure 2a and Supplementary Figure 2 for the brain features of each observer).

### Flow of Brain Features

For each observer, the full feature-by-voxel-by-time MI matrix comprises real-valued time series that represent the effect size for the representation of each brain feature in each voxel. In each observer, we noted that the representation of diagnostic and nondiagnostic brain features diverges around 170 ms post stimulus (see Supplementary Figure 2 for the full space by time representation of diagnostic and nondiagnostic brain features of each observer). On this basis, we defined an earlier (50-170ms post stimulus) and a later time window (170-400ms post stimuli) and summarized the representation of diagnostic and nondiagnostic brain feature in each. A voxel would represent diagnostic (vs. nondiagnostic) brain features if it has significant MI (FWER *p* < 0.05, one-tailed) for at least one diagnostic (vs. nondiagnostic) brain feature in this time window. For each voxel, we then counted the number of observers who satisfied these criteria and reported these distributions for diagnostic (white glass brains in Figure 2b) and nondiagnostic (magenta glass brains in Figure 2b) brain features, separately in the early and late time windows.

### Topology of Modal Brain Feature Representations

Using the full feature-by-voxel-by-time MI matrices of each observer (see above, *Method, Flow of Brain features*), within each time window we computed for each voxel the diagnostic or nondiagnostic brain feature(s) that was represented by a majority of observers (i.e. modally). A voxel is counted as representing a feature within a time window if its MI was significant for at least one time point. We performed this computation independently for each diagnostic brain feature of “the nuns,” or “Voltaire” and also for each nondiagnostic brain feature. Figure 2c reports the representation of these color-coded modal brain features, for voxel representing at least one diagnostic feature over N >=3 observers.

### Diagnostic and Nondiagnostic Brain Feature Representation in the Occipital-ventral pathway

For each individual observer, we proceeded in two steps:

#### Step1: Dynamics of diagnostic and nondiagnostic brain feature representation between 0 and 400ms post stimulus

For each observer, we used the full feature-by-voxel-by-time matrix computed above and selected the voxels with significant MI for at least one nondiagnostic brain feature in the 0 to 400ms time window (henceforth, “nondiagnostic voxels”). For each nondiagnostic voxel, we extracted at each time point the maximum MI over the nondiagnostic brain features (see Figure 3a for one typical observer, and Supplementary Figure 3-1 to 3-4 for the remaining observers). We defined a nondiagnostic voxel onset (vs. an offset) as the first (vs. last) time point at which the MI for any nondiagnostic feature is significant at this voxel (FWER *p* < 0.05, one-tailed), as well as its corresponding representation duration (computed as offset - onset). We further quantified, for each nondiagnostic voxel, their spatial organization by calculating the Euclidean Distance between each of these voxels and the voxel with the first significant onset for any nondiagnostic feature. We then performed the calculations just described for diagnostic voxels.

#### Step2: Locating the spatial-temporal junction of divergence between diagnostic and nondiagnostic feature representations

We selected the nondiagnostic voxels that represent nondiagnostic brain features furthest (i.e. with Euclidean distances > 75^th^ percentile of distances of all nondiagnostic voxels), and defined their maximum offset as the temporal marker of the junction (range between 155 and 177 ms across 5 observers, median = 163 ms).

To indicate the anatomical location of the junction voxels, we grouped nondiagnostic voxels (or the diagnostic voxels) of each observer according to their location in the cuneus (CU), lingual gyrus (LG), inferior occipital cortex (IOG), middle occipital gyrus (MOG), superior occipital gyrus (SOG), FG voxels locates quite close to LG (LG/FG, see Supplementary Figure 7 for location), fusiform gyrus (FG), inferior temporal gyrus (ITG), middle temporal gyrus (MTG), superior temporal gyrus (STG), inferior parietal lobe (IPL), and superial parietal lobe (SPL). In each anatomical region, we then mapped on the Y axis the Euclidean distance (see step 1) of all nondiagnostic voxels and all diagnostic voxels, cyan-yellow coded by their onset rank (see Figure 3c and Supplementary figures 3-1 to S3-4 for the results of other observers).

We defined temporal outlier voxels independently for diagnostic and nondiagnostic features, and excluded them from all above analyses. Outlier voxels were those > 3 standard deviations from the median onset of all voxels (computed separately with nondiagnostic and diagnostic voxel onset distributions, see Supplementary Table 2 for percentage of voxels exclusion).

### Feature Representation in the Brain for Perceptual Decision

For each individual observer, we proceeded in three steps:

#### Step 1. Computation of redundancy <Brain Feature Coefficients; MEG Voxel Activity; Perceptual Decision>

For each observer, brain voxel and diagnostic feature, we computed information theoretic redundancy, the triple relationship between <Brain Feature Coefficients; MEG Voxel Activity; “the nuns,“ “Voltaire,” “don’t know”>, every 2 ms between 112 and 244ms post stimulus. In each observer, we slightly adjusted the time boundaries to cover the first wave of processing encompassing the M/N100 and the M/N170 (as detailed in Figure 4b for one typical observer and Supplementary Figure 4-1 to 4-4 for the remaining observers). Redundancy measures the common variations between the relative presence of a diagnostic feature in the stimulus due to random sampling (cf. Figure 1), MEG voxel responses, and behavioral responses. We established statistical significance (FWER *p* < 0.05, one-tailed) by recomputing redundancy with shuffled decision responses across trials (repeated 200 times), and corrected for multiple comparison using the maximum statistics as explained above.

#### Step 2. Computation of feature representation complexity over each observer’s time window

For each observer, we constructed 5 evenly distributed time windows. In each, we summed the number of significant redundant features that each brain voxel represented. We used this sum to measure the complexity of redundant feature representation on each voxel and in each time window.

#### Step3. Extracting the redundant features for each perception

We classified each redundant feature in terms of their diagnosticity to each perception (see *Method, Diagnostic and Nondiagnostic Brain Features*). With diagnostic brain features of “the nuns” (and “Voltaire”), we calculated the complexity of redundant “nuns” (and “Voltaire”) feature representation on each voxel per time window as Step2

## Supplementary Information

**Supplementary Table 1.** Number of trials following pre-processing of MEG data.

| Observer | All responses | "Nuns" response | "Voltaire" response | "Don't Know" response |
| --- | --- | --- | --- | --- |
| 1 | 3314 | 1189 | 1313 | 812 |
| 2 | 3604 | 1666 | 1263 | 675 |
| 3 | 4154 | 1634 | 1892 | 628 |
| 4 | 3023 | 1603 | 738 | 682 |
| 5 | 2885 | 1007 | 1346 | 532 |

**Supplementary Table 2.** Percentage of voxels excluded from onset analysis.

| Observer | Diagnostic | Nondiagnostic |
| --- | --- | --- |
| 1 | 1.81% | 0.35% |
| 2 | 0 | 2.45% |
| 3 | 2.21% | 0.19% |
| 4 | 3.52% | 0 |
| 5 | 0.58% | 0 |

**Supplementary Figure 1.**
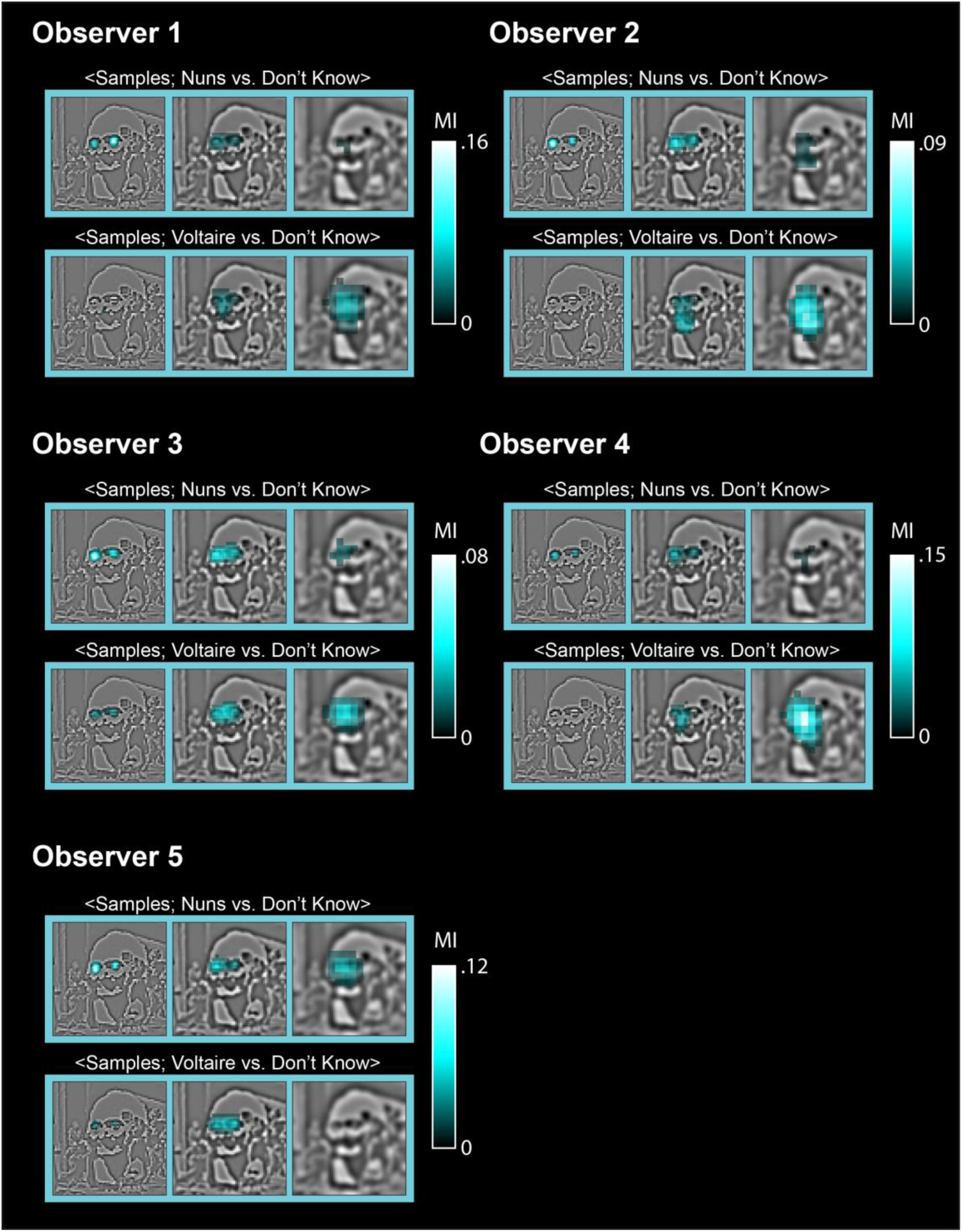
Diagnostic Features of Individual Observers. With Mutual Information, we quantified the single-trial relationship, for each sampled pixel in the 5 SF bands and separately for trials with responses <Information Samples; “nuns” vs. “don’t know”> and <Information Samples; “Voltaire” vs. “don’t know”>. The cyan framed images report the significant (FWER *p* < 0.01, one-tailed) pixels in the first three SF bands, revealing across observers the features most diagnostic of perceiving “the nuns” (the two small faces in SF band 1) and “Voltaire” (the broad face in SF band 3).

**Supplementary Figure 2.**
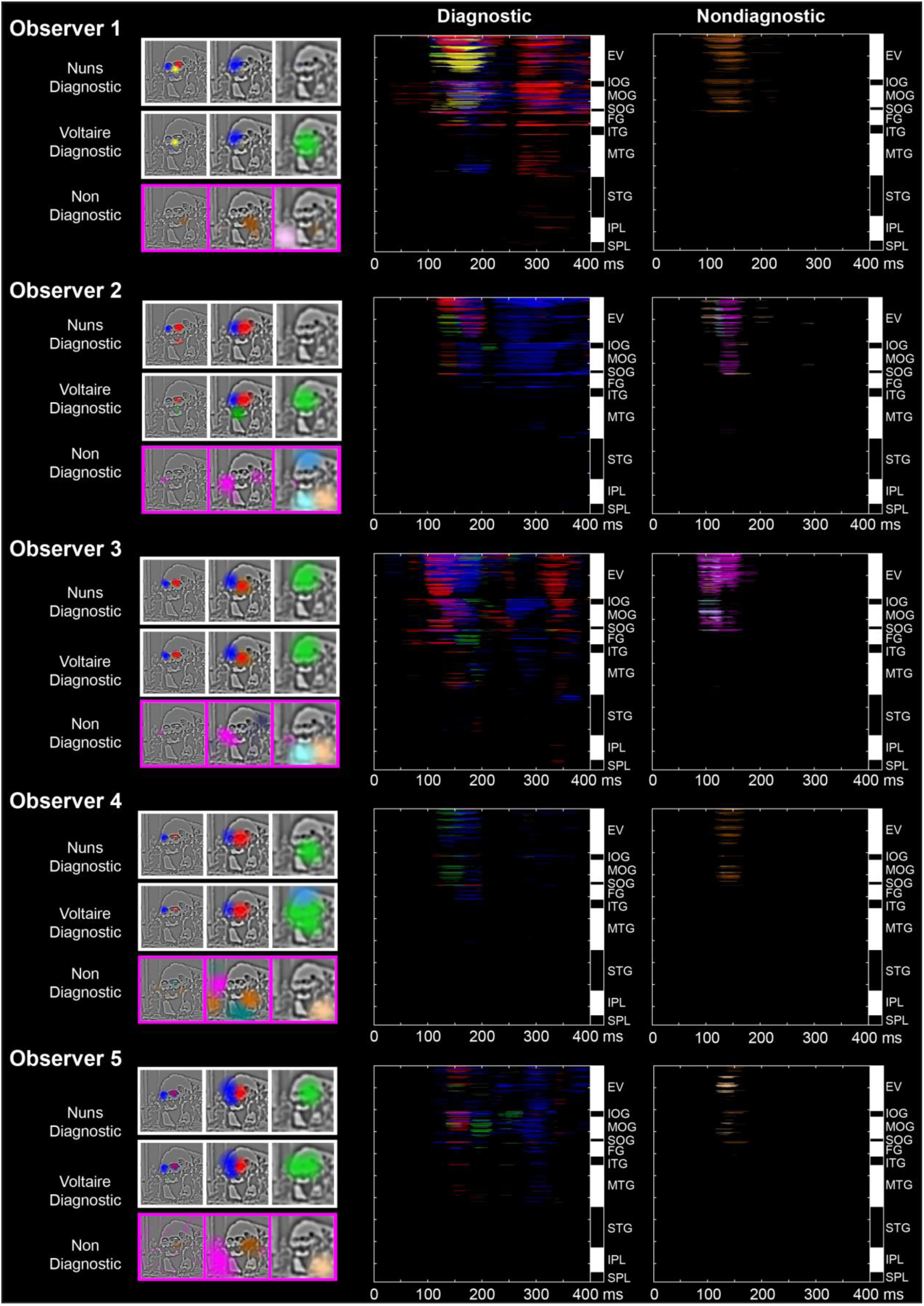
Representation of Diagnostic and Nondiagnostic Brain Features for Individual Observers. White frames highlight the diagnostic features that MEG voxels represent, separately presented for perceptual decisions of “the nuns” and “Voltaire.” Magenta frames highlight the features that MEG voxels also represent though they are not diagnostic of the perceptual decisions. Each cell of the Diagnostic and Nondiagnostic full brain feature-by-voxel-by-time matrices associated with each observer reports the color-coded brain feature with maximum significant MI across all brain features for this voxel and time point. The alternating white/black bars flanking the voxel axis of each matrix denote the anatomical brain region of the considered voxel. To illustrate with two examples, the matrices of Observer 1 reveal that the diagnostic brain feature “nose of Voltaire” in yellow is primarily represented in EV, MOG and FG across the full time course. In contrast, the nondiagnostic brain feature “to the right of Voltaire’s face” in brown is primarily represented in occipital regions, and before 180 ms. EV = early visual cortex (including lingual gyrus and cuneus), IOG = inferior occipital gyrus, MOG = middle occipital gyrus, SOG = superior occipital gyrus, FG = fusiform gyurs, ITG = inferior temporal gyrus, MTG = middle temporal gyrus, STG = superior temporal gyrus, IPL = inferior parietal lobe, SPL = superior parietal lobe.

**Supplementary Figure 3-1.**
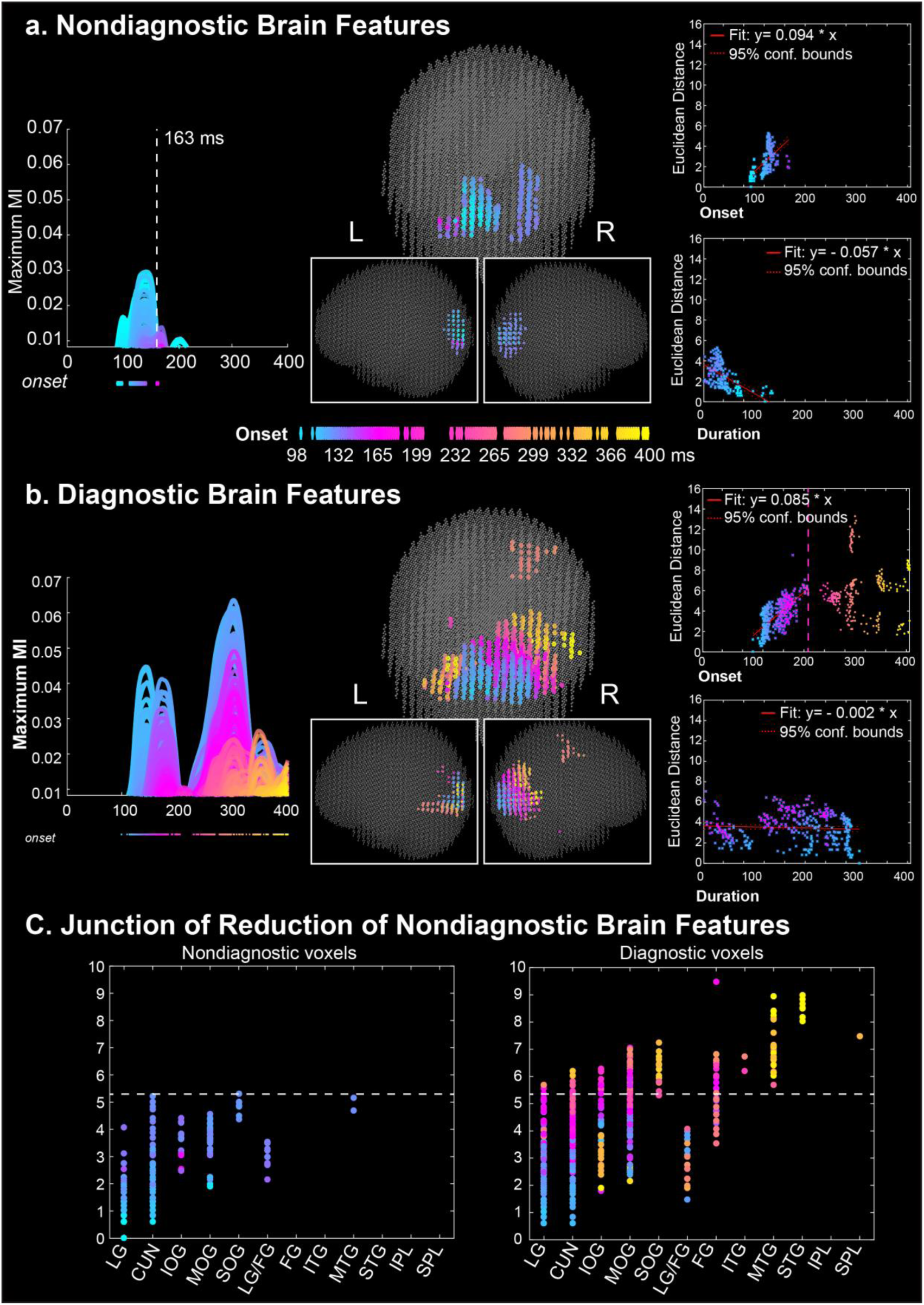
Reduction of Nondiagnostic Brain Feature Representations in Occipital Cortex. Organized as Figure 3, the results of observer 2.

**Supplementary Figure 3-2.**
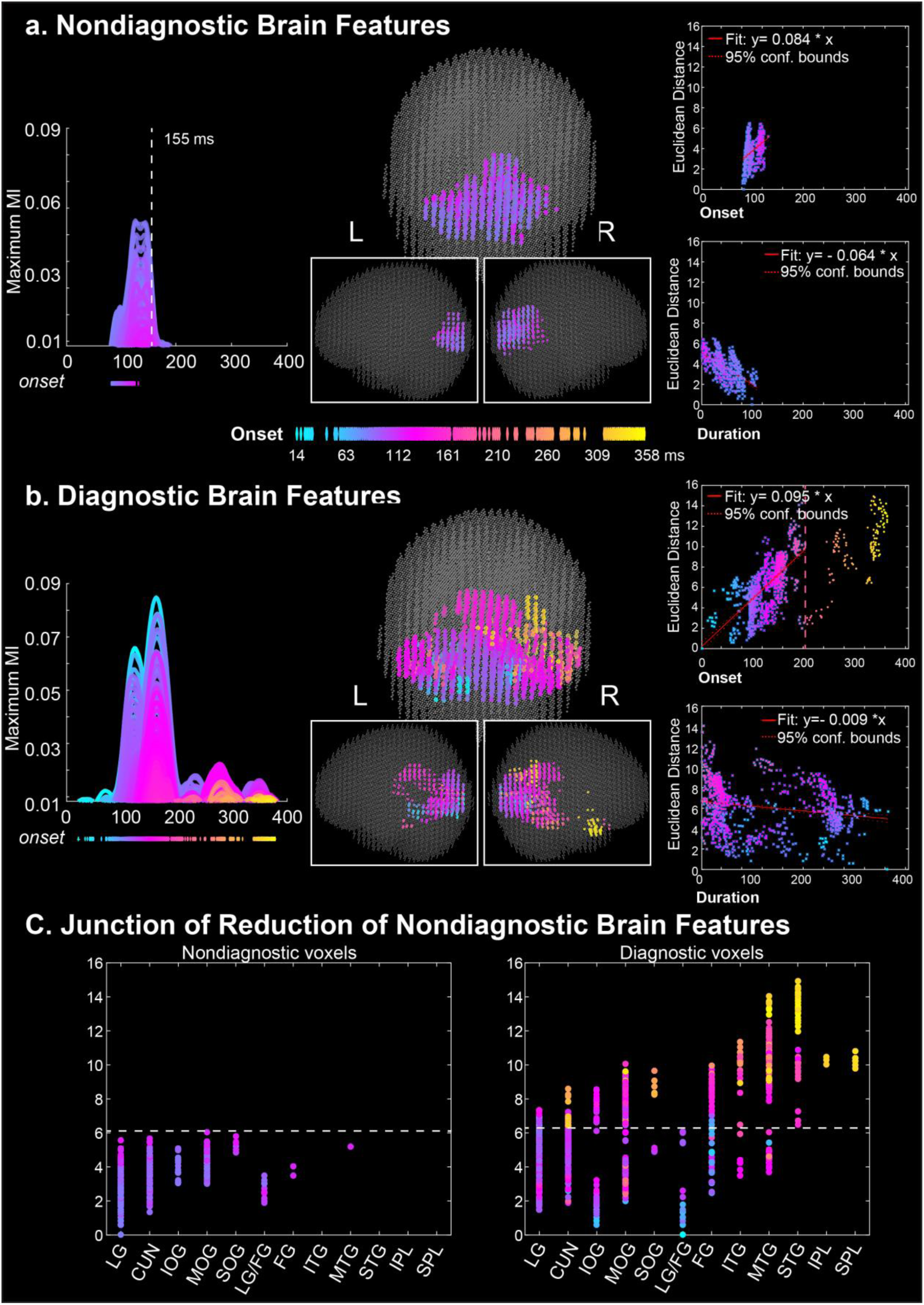
Reduction of Nondiagnostic Brain Feature Representations in Occipital Cortex. Organized as Figure 3, the results of observer 3.

**Supplementary Figure 3-3.**
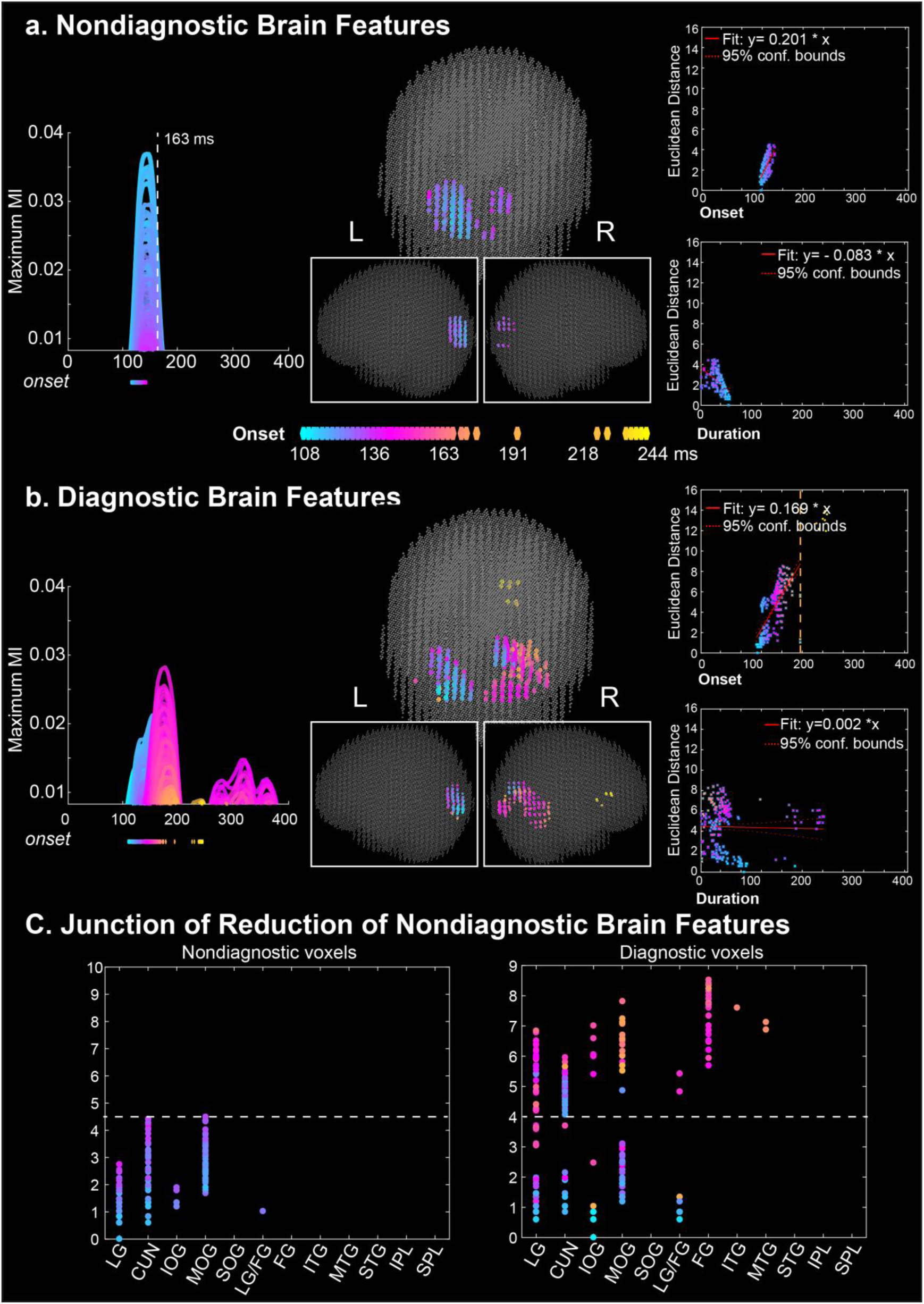
Reduction of Nondiagnostic Brain Feature Representations in Occipital Cortex. Organized as Figure 3, the results of observer 4.

**Supplementary Figure 3-4.**
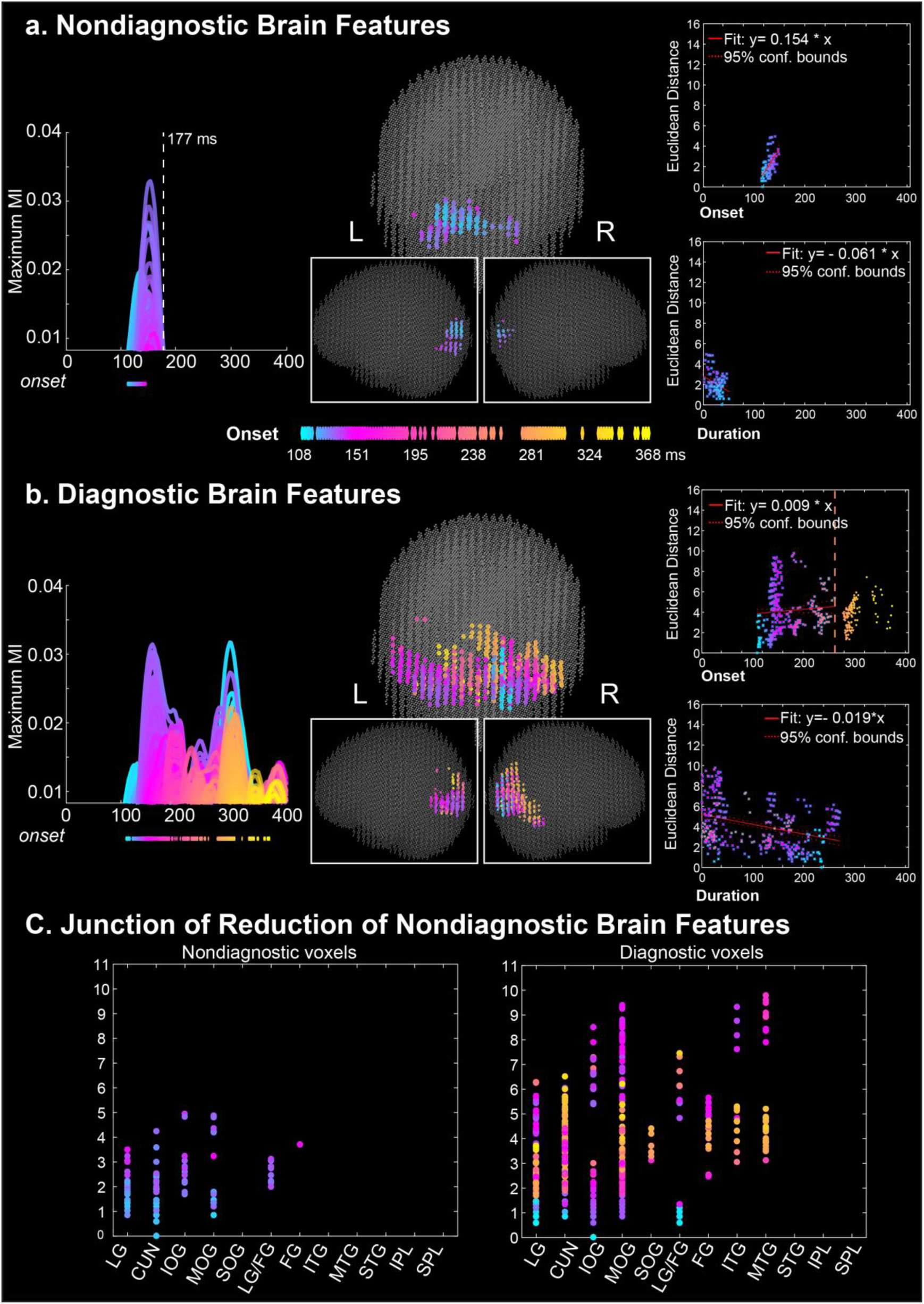
Reduction of Nondiagnostic Brain Feature Representations in Occipital Cortex. Organized as Figure 3, the results of observer 5.

**Supplementary Figure 4-1.**
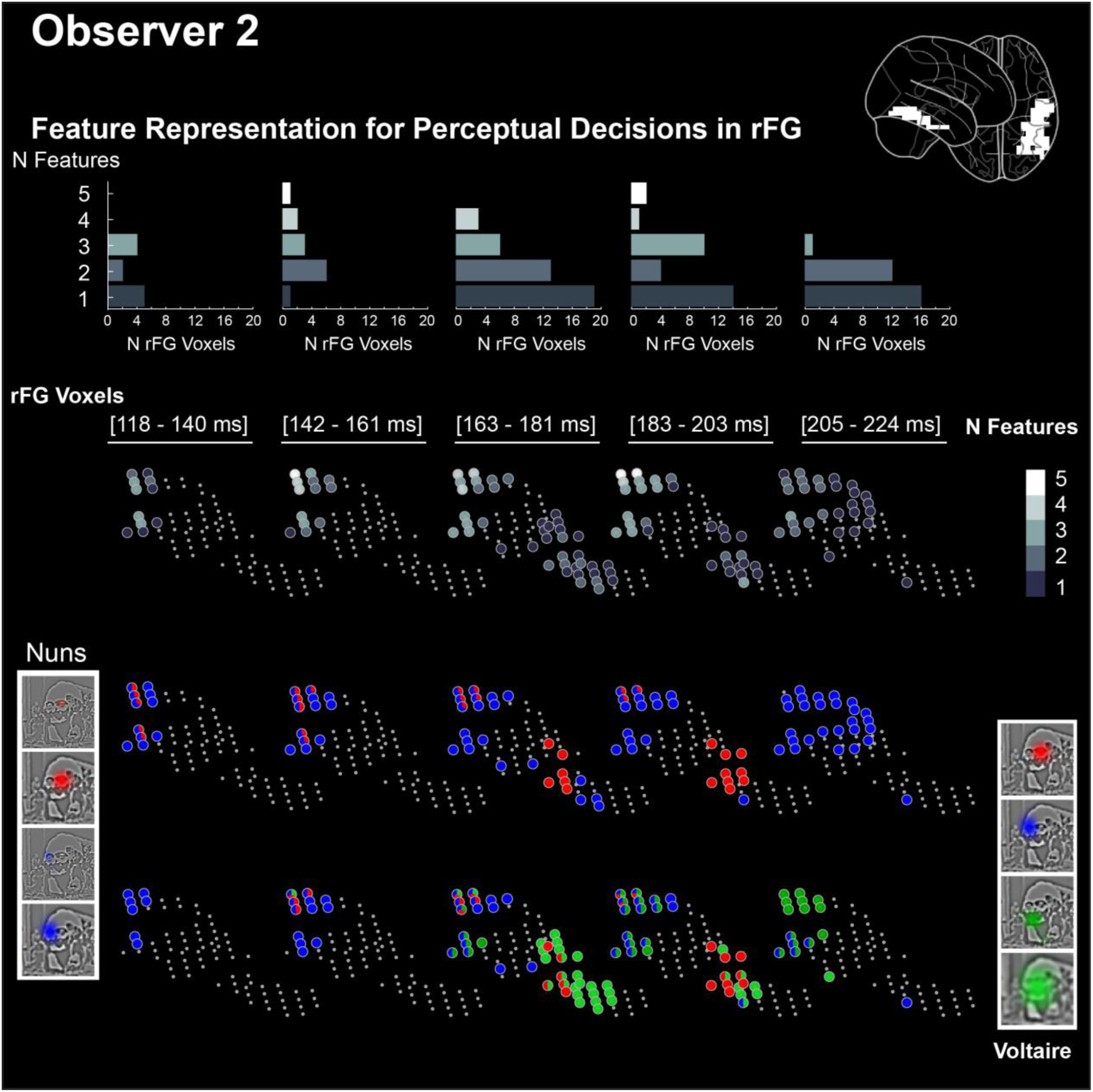
Dynamics of Perceptual Decision Representations in rFG. Organized as Figure 4, the results of observer 2.

**Supplementary Figure 4-2.**
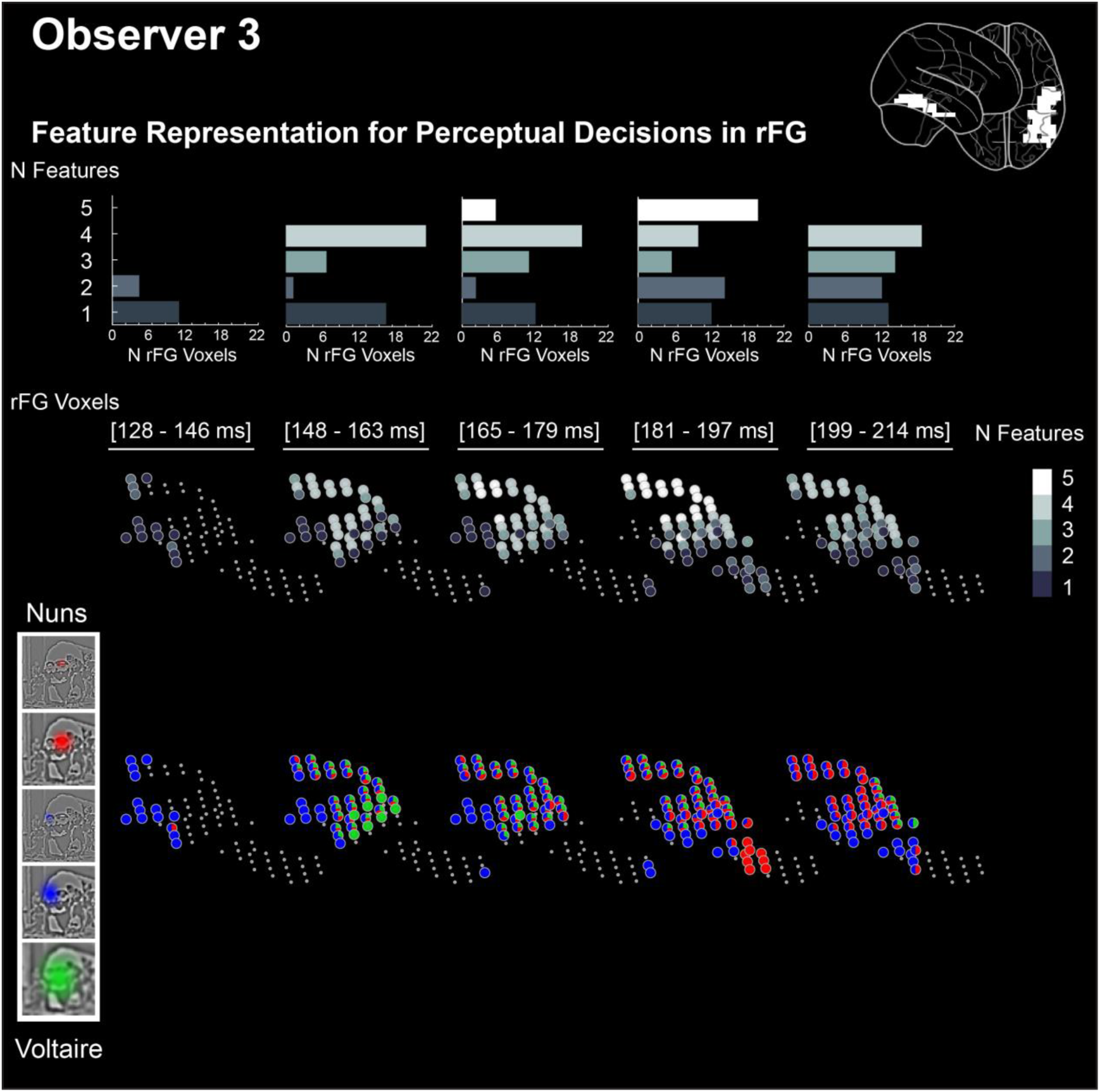
Dynamics of Perceptual Decision Representations in rFG. Organized as Figure 4, the results of observer 3.

**Supplementary Figure 4-3.**
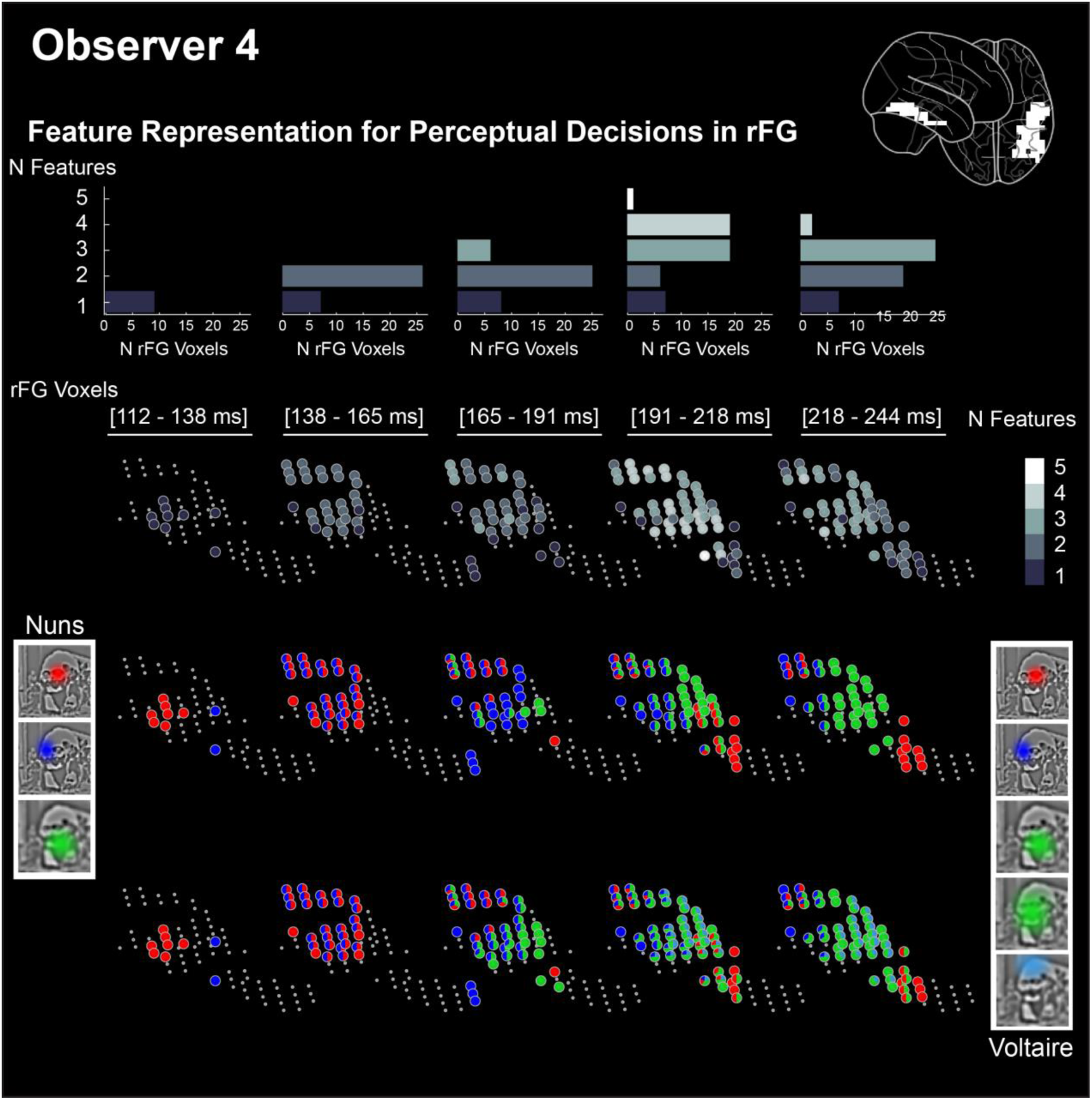
Dynamics of Perceptual Decision Representations in rFG. Organized as Figure 4, the results of observer 4.

**Supplementary Figure 4-4.**
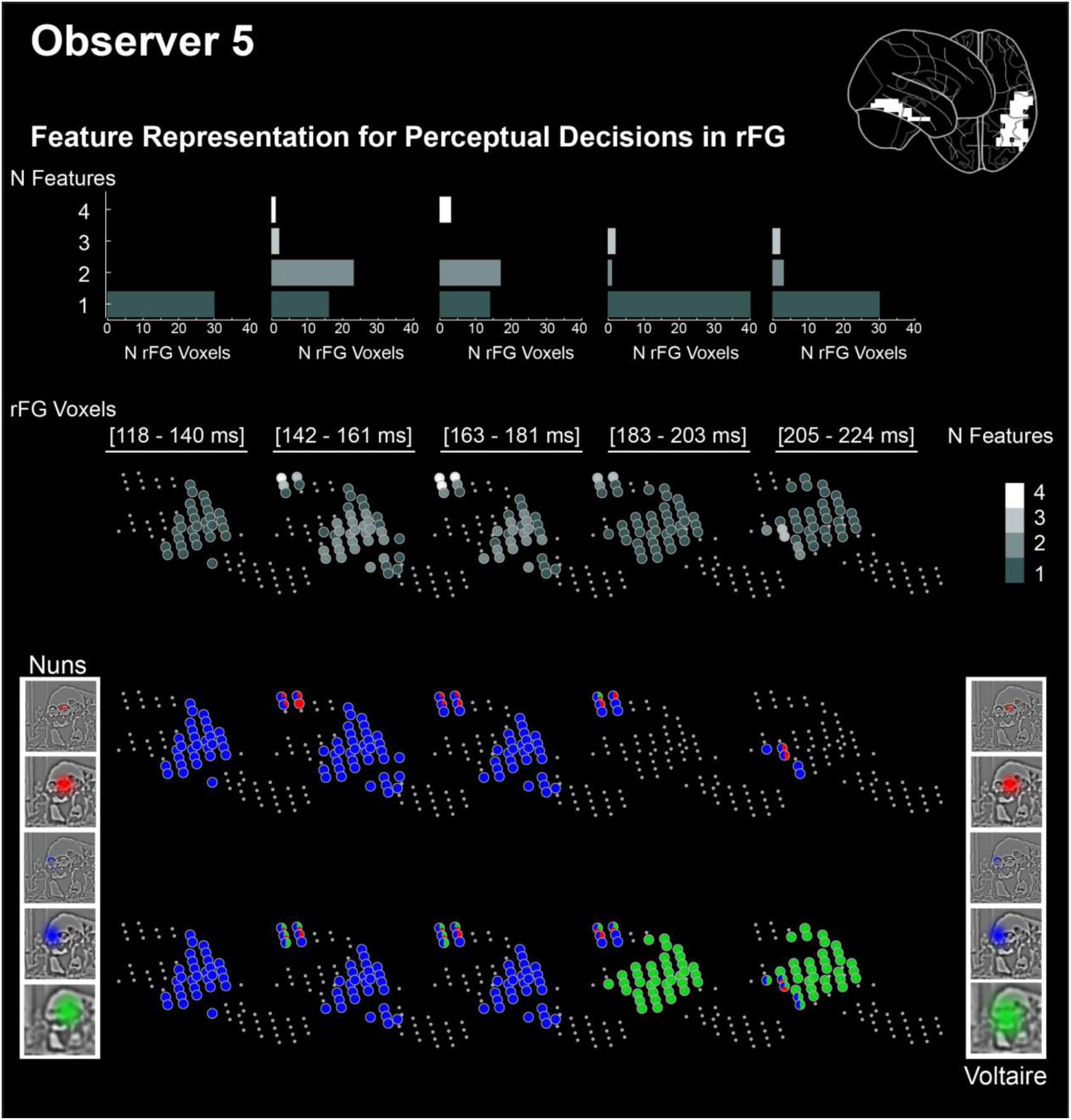
Dynamics of Perceptual Decision Representations in rFG. Organized as Figure 4, the results of observer 5.

**Supplementary Figure 5.**
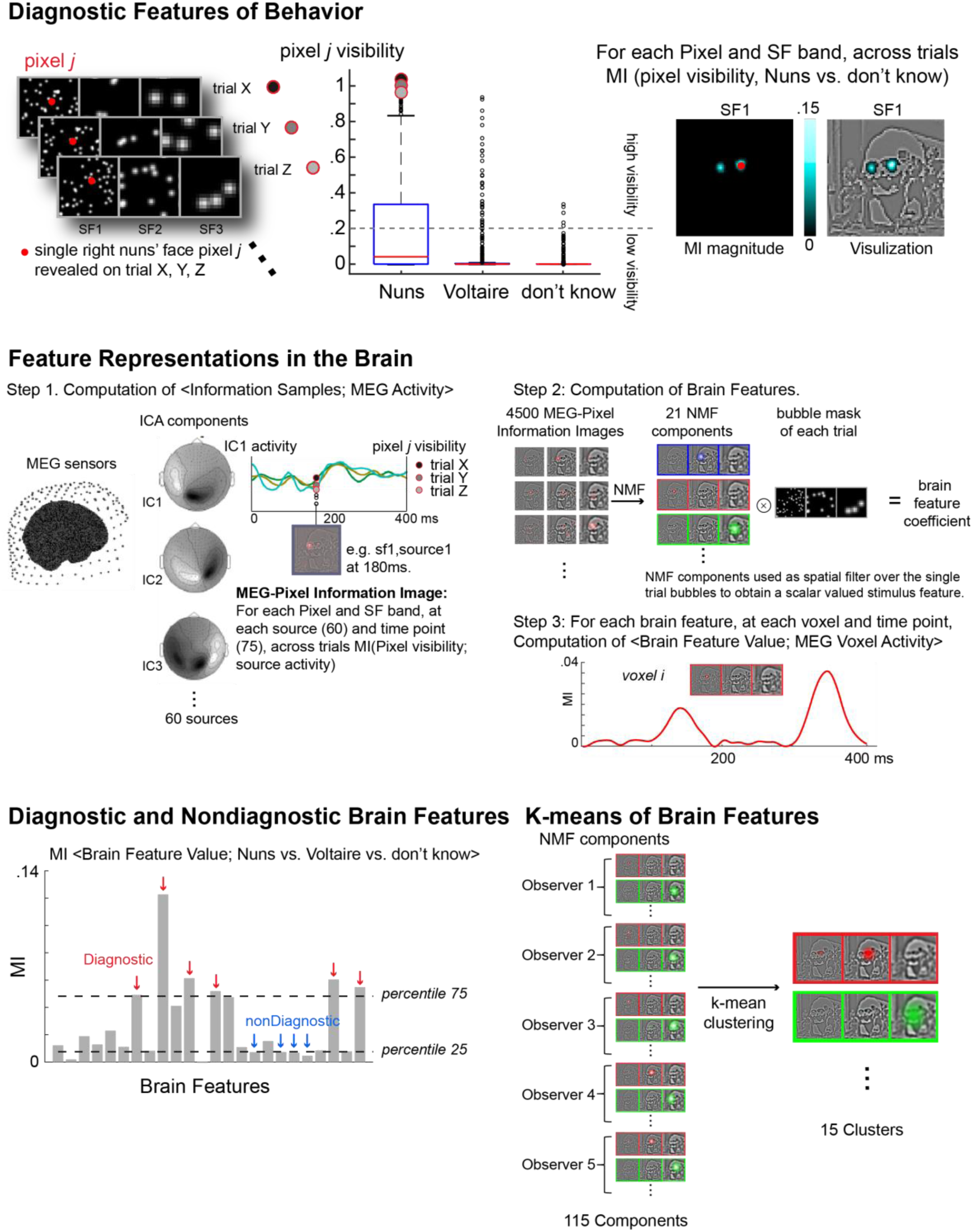
Details of the analysis pipeline. The *Diagnostic Features of Behavior* section shows example data for calculating the MI <Information Samples; “the nuns” vs. “don’t know”>. The *Feature Representations in the Brain* section illustrates the process to obtain Brain Features (step1&2) and the calculation of MI <Brain Features; MEG Voxel Activity> (step 3). The *Diagnostic and Nondiagnostic Brain Features* section illustrate how we use the MI distribution <Brain Feature Value of Sampled Stimuli; Perceptual decision> to determine the types of brain features. The *K-means of Brain Features* section uses example data to illustrate the k-means clustering analysis.

**Supplementary Figure 6.**
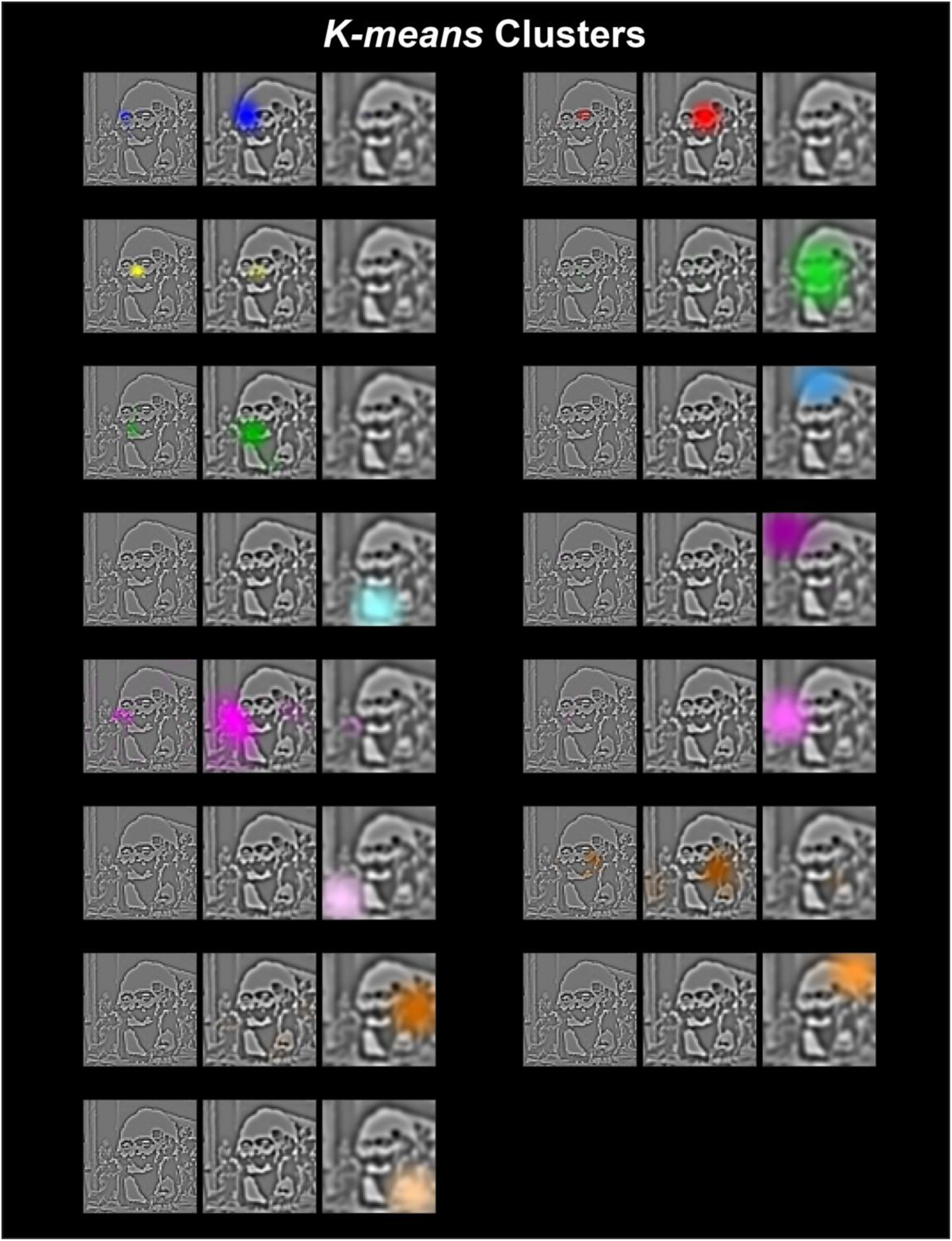
The output features of k-means clustering on the brain features of 5 observers. We color-coded each feature.

**Supplementary Figure 7.**
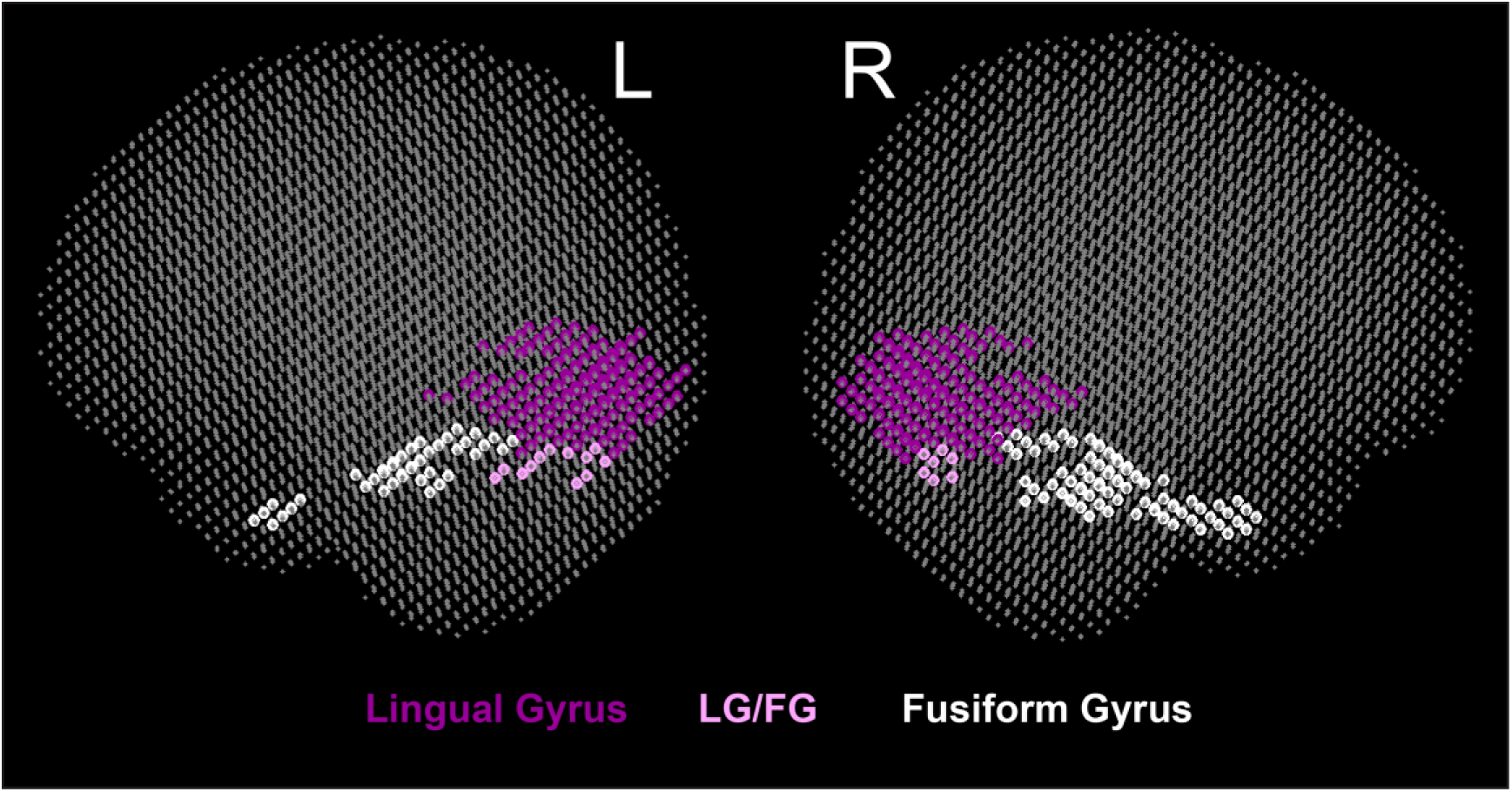
Location of LG/FG voxels. The dark purple scatters show lingual gyrus (LG) voxels; the light purple scatters show LG/FG voxels which are fusiform gyrus voxels located next to lingual gyrus voxels; the white scatters show the FG voxels we included in *Feature Representation for Perceptual Decision* analysis.

